# Kinetic pathway of ATP-induced DNA interactions of ParA2, a protein essential for segregation of *Vibrio cholerae* chromosome 2

**DOI:** 10.1101/2021.02.27.433207

**Authors:** Satpal S. Chodha, Adam C. Brooks, Peter Davis, Revathy Ramachandran, Dhruba K Chattoraj, Ling Chin Hwang

**Affiliations:** Department of Molecular Biology and Biotechnology, University of Sheffield, Firth Court, Western Bank, Sheffield, United Kingdom; Basic Research Laboratory, Center for Cancer Research, National Cancer Institute, National Institutes of Health, Bethesda, MD, United States

**Keywords:** chromosome segregation, plasmid partition, ParA ATPase, diffusion-ratchet, dynamic oscillations

## Abstract

*Vibrio cholerae* chromosome 2 (Chr2) requires its own ParABS system for segregation. Without it, *V. cholerae* becomes nonviable and loses pathogenicity. ParA2 of Chr2 is a Walker-type ATPase that is the main driver of Chr2 segregation. Most of our understanding of ParA function comes from studying plasmid partition systems. How ParA provides the motive force in segregation of chromosomes, which are much larger than plasmids, is less understood and different models have been proposed. Here we analyzed *in vivo* behavior and kinetic properties of ParA2 using cell imaging, biochemical and biophysical approaches. ParA2 formed an asymmetric gradient in the cell that localized dynamically in the cell cycle. We found that ParA2 dimers bind ATP and undergo a slow conformational change to an active DNA-binding state, similar to P1 ParA. The presence of DNA catalyzes ParA2 conformational change to allow cooperative binding of active ParA2 dimers to form higher-order oligomers on DNA. Nucleotide exchange rates were also slow, thus providing a control of ParA2 recruitment and dynamic localizations. Although highly conserved in biochemical properties, ParA2 showed faster overall ATP cycling and DNA-rebinding rates than plasmid ParAs, suggesting that this could be shared kinetic features among chromosomal ParAs to regulate the transport of a much larger DNA cargo.

## INTRODUCTION

Chromosome replication and segregation are fundamental cellular processes in all kingdoms of life. Replicated DNA must be accurately distributed to progeny cells before cell division. Eukaryotic cells employ the mitotic spindle apparatus for the segregation of sister chromatids. In contrast, most bacteria use the tripartite ParABS system to actively segregate their replicated chromosomes to progeny cells (Baxter & Funnell, 2015; Gerdes *et al*, 2004; Reyes-Lamothe *et al*, 2012). The ParABS system consists of centromere-analogous *parS* sequences, a *parS*-binding protein ParB, and a ParA ATPase that binds to DNA nonspecifically. The chromosomal Par proteins also contribute to other key cellular processes including DNA condensation, chromosome replication, transcription regulation, cell division and motility (Bouet *et al*, 2014; Baek *et al*, 2014; Jha *et al*, 2012; Lasocki *et al*, 2007). The Par system is essential for the survival and pathogenesis of bacteria with multipartite genomes such as *Vibrio cholerae* and *Burkholderia cenocepacia* (Yamaichi *et al*, 2007b; Du *et al*, 2016).

*V. cholerae*, the causative agent for cholera, has two chromosomes: a larger 3 Mb chromosome 1 (Chr1) and a smaller 1 Mb chromosome 2 (Chr2). Both chromosomes encode ParABS systems that are specific to their own segregation, which display distinct and independent dynamics (Jha *et al*, 2012; Fogel & Waldor, 2006; Ramachandran *et al*, 2014). Upon replication, one of the sister Chr1 origins travels asymmetrically from the old pole to the new pole, while Chr2 origin relocates symmetrically from the mid-cell to quarter-cell positions (Fogel & Waldor, 2006; Srivastava & Chattoraj, 2007). The migration of the replication origins of Chr1 and Chr2 are directed by the partition complexes, which consists of ParB forming foci with the cognate *parS* sites close to the origin region (Yamaichi *et al*, 2007a). The motion and segregation of the partition complex is driven by interactions with ParA. ParA1 of Chr1 forms a retracting cloud moving ahead of the ParB1 foci toward the new cell pole (Fogel & Waldor, 2006; Kadoya *et al*, 2011). It was proposed that Chr1 uses a mitotic-like mechanism where ParA1 filaments attach to ParB1 loci and depolymerize to pull the sister loci apart (Fogel & Waldor, 2006). The ParABS1 system is not essential for the segregation of Chr1, which apparently employs an as yet unknown mechanism for segregation in absence of the Par system (Yamaichi *et al*, 2007a). On the contrary, the ParABS2 system is essential for Chr2 segregation and *V. cholerae* viability. Deletion of *parAB2* loci results in loss of Chr2 over time and activation of toxin-antitoxin systems that kills cells, thereby causing loss of *V. cholerae* pathogenesis (Yamaichi *et al*, 2007b). Despite the vital role ParA2 plays in *V. cholerae* proliferation, very little is known about the behavior and biochemistry of Chr2 segregation.

Most of our understanding of the ParABS apparatus stems from studies of plasmid partition systems. The systems are categorized according to the type of ParA they have. Type II and Type III ParAs are actin- and tubulin-like NTPases, respectively, as they are biochemically and structurally similar to their eukaryotic homologues (Gerdes *et al*, 2004; Baxter & Funnell, 2014; Ebersbach & Gerdes, 2005). Similarly, their plasmid partition systems employ filament-based mechanisms to polymerize and generate forces that push or pull plasmids apart (Gerdes *et al*, 2010; Szardenings *et al*, 2011). However, most bacterial chromosomes including *V. cholerae* and low copy number plasmids use the ubiquitous Type I deviant Walker-type ATPase for DNA segregation (Reyes-Lamothe *et al*, 2012; Bouet *et al*, 2014; Ramachandran *et al*, 2014). Numerous biochemical studies have been done on ParAs from plasmids, such as P1, F, TP228, and pB171 and pSM19035 (Davey & Funnell, 1997; Bouet *et al*, 2007; Ah-Seng *et al*, 2009; Vecchiarelli *et al*, 2010; Soberón *et al*, 2011; Pratto *et al*, 2008; Barillà *et al*, 2007, 2005; Bouet & Funnell, 1999; Davis *et al*, 1992, 1996; reviewed in Baxter & Funnell, 2014). In general, ParA dimerizes upon ATP binding and has a weak ATPase activity that is stimulated by its cognate partner ParB and non-specific DNA. ParA dimers have a crucial nonspecific DNA binding activity that allows them to associate with (chromosomal) nucleoid DNA, which is exploited as a mechanism for moving and segregating plasmids across the cell (Vecchiarelli *et al*, 2010, 2012). As demonstrated by P1 ParA, a critical feature in the DNA binding step is the slow conformational change of ParA upon ATP-binding, from a freely diffusing state to a DNA-binding state (Vecchiarelli *et al*, 2010). Based on *in vitro* reconstitution of P1 and F plasmid partition dynamics, a brownian/diffusion-ratchet mechanism was proposed. The slow rebinding of ParA on DNA relative to its fast disassembly by ParB-*parS* plasmids generates a ParA concentration gradient on the nucleoid, driving plasmid dynamics. (Hwang *et al*, 2013; Vecchiarelli *et al*, 2013; reviewed in Brooks & Hwang, 2017). The moving ParA gradient drive directed motion of the plasmid via successive ParA-ParB interactions toward higher ParA concentrations at cell poles (Vecchiarelli *et al*, 2014; Hu *et al*, 2015, 2017). This diffusion-ratchet mechanism has little similarity to the mitotic type mechanism proposed for Chr1 segregation.

A modification of the diffusion-ratchet mechanism has been subsequently proposed in *C. crescentus* where the partition complex is ‘relayed’ across the cell by utilizing the intrinsic elastic properties of the chromosome (Lim *et al*, 2014; Surovtsev *et al*, 2016). F SopA and *B. subtilis* Soj were found to localize within the high density regions of the nucleoid, which could allow plasmids to ‘hitchhike’ along those regions (Le Gall *et al*, 2016). An oscillating TP228 ParF meshwork that permeates the nucleoid has been observed and proposed to ‘trap’ and release plasmids to localize them (McLeod *et al*, 2017; Caccamo *et al*, 2020). These and other cellular studies that exhibit ParA gradients have built their models on either the diffusion-ratchet or filament-based mechanism (Fogel & Waldor, 2006; Hu *et al*, 2015; Lim *et al*, 2014; McLeod *et al*, 2017; Ptacin *et al*, 2010; Walter *et al*, 2017; Hatano & Niki, 2010; Hatano *et al*, 2007; Ringgaard *et al*, 2009; Jindal & Emberly, 2015). However, the key parameter – the time delay from non-binding to DNA-binding state that allows for ParA gradient formation – has only been shown for P1 plasmid (Vecchiarelli *et al*, 2010). It is still unknown if any other plasmid or chromosome ParAs share similar or different biochemical properties that license them to spatiotemporally pattern the nucleoid for DNA segregation. A key question remains as to whether (and how) similar ParA gradients that suffice for plasmids, could translocate the chromosome, a much larger DNA cargo, across the cell.

To understand the molecular basis of ParA-mediated chromosome segregation, we first performed *in vivo* imaging of *V. cholerae* ParA2 and observed that it formed gradients that displayed dynamic pole-to-pole localizations. We then performed biochemical and biophysical characterizations of the kinetics of ParA2 ATPase cycle and DNA binding. We found that ParA2 undergoes a slow conformational change and slow ADP to ATP exchange step that are involved in ParA2 gradient formation and dynamics, indicating diffusion-ratchet mechanism to be the basis for Chr2 segregation. The reaction rates of ParA2 were faster than those of plasmid ParAs, implying that ParA2 has evolved to be a more robust and efficient enzyme, which might be required to regulate the transport of a much larger DNA cargo and to be more responsive in coordinating chromosome segregation dynamics to the timing of cell division.

## RESULTS

### ParA2 display dynamic localization in *V. cholerae*

To monitor the spatiotemporal distribution of ParA2 in wild type *V. cholerae* cells, we expressed ParA2-GFP from a plasmid encoding ParA2 with a C-terminal GFP fusion. The presence of the plasmid did not alter the generation time or cell-length distribution significantly, nor did chubby cells appear when Chr2 is lost (Yamaichi *et al*, 2007b). We observed that almost all of the cells that expressed ParA2-GFP had an asymmetric distribution of the protein along the long-axis of the cell (98%, n=61) (Figure 1A). The distribution showed a comet-like gradient with the highest intensity toward one of the cell poles that gradually decreased as it approached the cell center in predivisional cells, or to the septum in dividing cells. In time-lapse imaging, the ParA2-GFP gradients showed variability in the spatiotemporal dynamics. 51% of the cell population showed ParA2-GFP gradients either persisting at the same location or migrating to the opposite pole once in the same cell cycle (Figure 1A). On the other hand, 49% of the cells showed more dynamic localizations of ParA2-GFP, a typical example of which can be seen in Figure 1B. In these cells, ParA2-GFP gradients started from one pole and transitioned to form the gradient at the opposite pole before diffusing back again. Several periods of pole-to-pole oscillation occurred during each cell cycle. The oscillation periods varied from 5 to 6 min and pole-to-pole transition times from 30 to 60 s. The residence times of ParA2-GFP at the poles were longer. In some incipient mother cells, ParA2-GFP gradients were seen at both the cell poles. In these cells, sometimes the ParA2-GFP began to oscillate between the pole and the closing septa.

**Figure 1.**
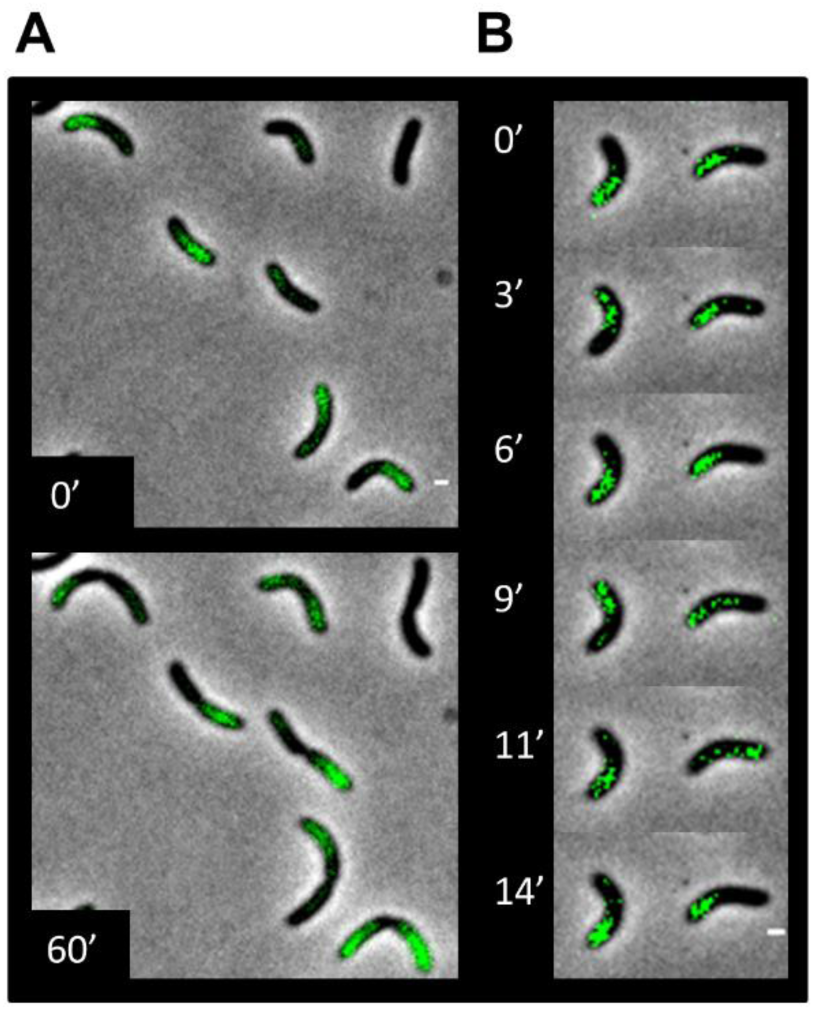
Spatiotemporal localization of ParA2-GFP in *V. cholerae* cell culture. Representative cells carrying plasmids expressing ParA2-GFP. **(A)** A population of cells displaying decreasing gradients of ParA2-GFP from a cell pole at 0 min (top) and at 60 min (bottom). **(B)** Time-lapse imaging of cells showing pole-to-pole oscillations of ParA2-GFP gradients. The scale bar is 1 μm.

Dynamic bands of *Vc* ParA1, belonging to Chr1 was reported earlier, which appears distinct from ParA2 (Fogel & Waldor, 2006). ParA1 forms a tight focus at one (old) pole and a dense band at the opposite (new) pole that retracts toward the new pole once per cell cycle, with ParB1 foci trailing. ParA1 retraction has been interpreted as the mitotic-like pulling of ParA1 polymers (Fogel & Waldor, 2006). Similar oscillatory dynamics to ParA2 have also been reported for ParA homologs in plasmids and chromosomes (McLeod *et al*, 2017; Hatano *et al*, 2007; Ringgaard *et al*, 2009; Marston & Errington, 1999; Ah-Seng *et al*, 2013). To understand how might ParA2 spatiotemporal patterns form and mediate Chr2 segregation, we next characterized the biochemical and kinetic properties of *Vc* ParA2.

### Apo ParA2 dimerize without nucleotides

ParA2 belongs to the family of P-loop ATPases that contains a deviant Walker A motif (**K**GGTG**K**S) that is involved in ATP-binding and hydrolysis. It was previously determined that ParA2 does not form higher-order polymers with ATP (Hui *et al*, 2010). Instead, our results on size exclusion chromatography multi-angle light scattering (SEC-MALS) analysis of ParA2 at 40 μM showed that the majority of the population formed dimers with an average MW of 91.1 kDa (Figure 2A) (theoretical MW 92.8 kDa). The peak positions and widths of elution profiles remained relatively unchanged in the presence of ADP and ATP. The sample polydispersity index remained very low with a mean of 1.014 across all samples. However, the peak MW remained slightly below that expected from a dimer, with a mean equivalent mass of 1.89 monomers. This lower than expected mass is most likely the result of a very small population of monomeric species present in equilibrium with dimers, regardless of the presence of nucleotide. This is in contrast with P1 ParA, which when analyzed at similar concentrations, is in a monomer-dimer equilibrium without nucleotide and stabilizes as a dimer with ATP or ADP (Vecchiarelli *et al*, 2010).

**Figure 2.**
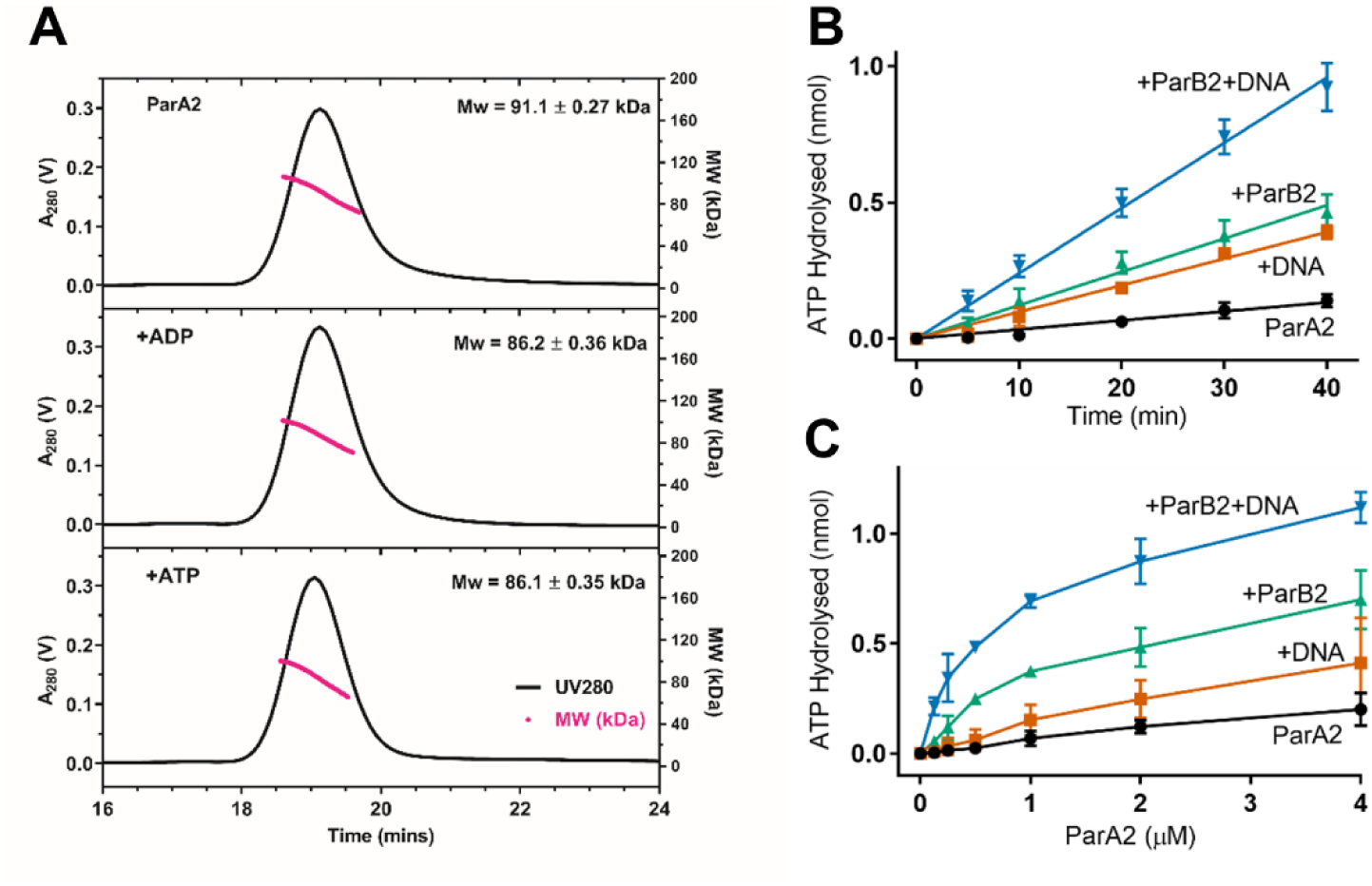
Association state and ATPase activity of ParA2. **(A)** ParA2 dimerizes in absence and presence of adenine nucleotides. SEC/MALS profiles of ParA2 in absence of nucleotide (top), in the presence of ADP (middle) and ATP (bottom). 40 µM ParA2 was pre-incubated with 2 mM nucleotide for 15 min, when present. Running buffer contained 0.5 mM nucleotide. **(B)** ParA2 pre-steady state ATP hydrolysis kinetics. 1.5 µM ParA2 was mixed in buffer A with 200 µM ATP spiked with 64 nM [α-^32^P]-ATP. 1.5 µM ParB2 and 100 µg ml^−1^ sonicated salmon sperm DNA were added where indicated. The hydrolysis products were measured at 23°C after the indicated reaction times. **(C)** ParA2 steady state ATP hydrolysis. Same as in (B), except that ParA2 concentration was varied as indicated and hydrolysis products were measured after a fixed interval of 30 min.

### ParA2 ATPase activity is stimulated by ParB2 and DNA

Next, we measured the ATPase activity of ParA2. We incubated ParA2 with [α^32^P]-ATP and used thin layer chromatography to separate the hydrolyzed products. We found that ParA2 is a weak ATPase with specific activity 0.1 mol ATP min^−1^mol^−1^ that is 2-fold higher than those of P1 ParA and F SopA (Figure S1). Similar to plasmid ParAs, its pre-steady state ATP hydrolysis rate was stimulated 2-fold by nonspecific DNA and 3-fold by its cognate partner *Vc* ParB2 (Figure 2B). The stimulation increased to 8-fold in the presence of both DNA and ParB2. Steady state ATPase activity of increasing ParA2 concentrations showed these pre-steady state rates were close to maximum under these conditions (Figure 2C). The amount of ATP hydrolyzed saturated towards higher ParA2 concentrations due to depletion of [α^32^P]-ATP substrate (Figure 2C). In summary, ParA2 ATPase activity is stimulated by both DNA and ParB2, similar to plasmid ParAs, but quantitatively is higher for *Vc* ParA2.

### Investigation of nucleotide binding and exchange rates

P1 ParA was found to undergo a slow multi-step conformational change upon ATP binding that was proposed to generate an uneven distribution of the protein on the nucleoid, required for plasmid motility (Vecchiarelli *et al*, 2010). To examine if *V. cholerae* uses a similar mechanism for ParA2 localization and Chr2 segregation, we sought to find the rate-limiting-step in the ATPase cycle of ParA2. Using fluorescently-labeled adenine nucleotides, MANT-ATP and MANT-ADP, we determined their binding affinities to ParA2. At steady state, ParA2 bound MANT-ATP and MANT-ADP similarly with *K_d_* = ∼11 μM (Figure S2). This affinity is 2- to 9-fold higher than that reported for *Vc* ParA2: 22 μM (ATP) and 34 μM (ADP) (Hui *et al*, 2010) and all other ParA homologs: P1 ParA 30 μM (ATP) and 50 μM μM (ADP) (Davey & Funnell, 1997; Vecchiarelli *et al*, 2010), F SopA 74 μM (ATP) (Bouet *et al*, 2007), *C. crescentus* ParA 50-60 μM (ATP) (Easter & Gober, 2002) and TP228 ParF 100 μM (ATP) (Barillà *et al*, 2005). ParA2 thus has higher binding affinities to ATP and ADP than other characterized ParA proteins.

We next monitored the fast kinetics of MANT-labeled nucleotides interactions with ParA2 using a stopped-flow fluorimeter. The relative fluorescence of MANT-ATP increased with binding to ParA2 over time and the extent of binding scaled proportionately with ParA2 concentrations (Figure 3A). The binding curves fitted well to a single exponential association with time constants, τ = 7-12 s for MANT-ATP and took almost 30 s to reach an apparent steady state. This timescale of binding ATP appears to be slow for a typical enzyme, suggesting that ParA2 may be undergoing dimerization or protein conformational change with ATP-binding. However, at higher ParA2 concentrations, the observed MANT-ATP association rates did not increase significantly (Table S1). This suggests that ParA2 already pre-exists as dimers at concentrations from 0.6 μM, prior to binding ATP, this was also reflected in the single-exponential fits. The pseudo-first-order rate constants, k_on_ and k_off_, of MANT-ATP binding to ParA2 were also determined by titrations of increasing MANT-ATP concentrations (Figure 3B). MANT-ADP association rates were also similar to those of MANT-ATP and remained relatively constant with increasing ParA2 concentrations (Figure 3C and Table S1). k_on_ and k_off_ of MANT-ADP were similar to MANT-ATP, and their calculated *K_d_*s (k_off_/k_on_) were both ∼8 μM, similar to that determined from steady state measurements (Figure S2). Next, we examined the dissociation rates of MANT-labeled nucleotides from ParA2 by preincubating ParA2 with MANT-AXP before mixing with unlabeled nucleotides. The subsequent decrease in MANT-ATP fluorescence was multiphasic, with an initial fast phase followed by a slower phase of τ = 50-60 s (Figure 3E), indicating that ATP-hydrolysis slows down nucleotide release from ParA2. In comparison, MANT-ADP fluorescence dropped rapidly within τ = 12-14 s, almost 5-fold faster than MANT-ATP (Table S1). In sum, it appears that ADP binds ParA2 similarly as ATP but dissociates more easily than ATP.

**Figure 3.**
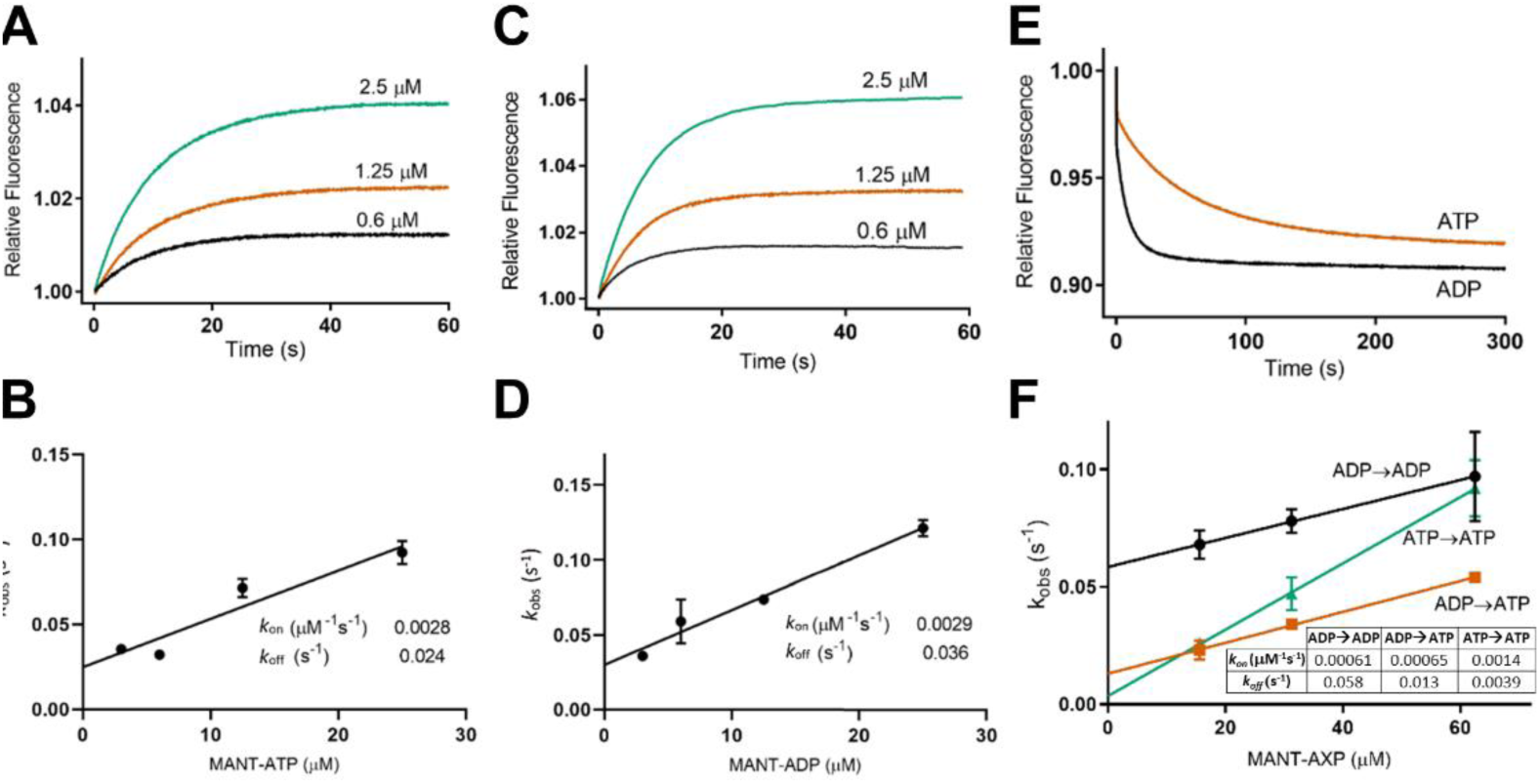
(A) ParA2-MANT-ATP binding kinetics. ParA2 at indicated concentrations and 25 μM MANT-ATP were prepared separately in Buffer B. Stopped-flow fluorescence spectroscopy was used to mix rapidly and monitor the change in relative MANT fluorescence. **(B)** Plot of pseudo-first order rate constant *k_obs_* against MANT-ATP concentration. Samples were prepared as in (A), except MANT-ATP was 10x higher concentration than ParA2. **(C)** ParA2-MANT-ADP binding kinetics. As in (A), except with MANT-ADP. **(D)** Plot of pseudo-first order rate constant *k_obs_* against MANT-ADP concentration. Samples were prepared as in (C), except MANT-ADP was 10x higher concentration than ParA2. **(E)** ParA2-MANT-AXP dissociation kinetics. ParA2, at indicated concentrations, was prepared with MANT-AXP in a 1:2 ratio, while 1 mM AXP was prepared separately. ParA2 and MANT-AXP were pre-incubated at 23°C for 3 min, then rapidly mixed with AXP. Dissociation kinetics were subsequently measured as a decrease in relative MANT fluorescence. **(F)** ParA2-AXP exchange kinetics plot. *k_obs_* plotted against MANT-AXP concentration. ParA2 and AXP were prepared to indicated concentrations, in a 1:5 ratio, while MANT-AXP was prepared separately and at a 5x higher concentration than AXP.

In growing cells, ParA2 dimers would undergo ADP→ATP nucleotide exchange upon ATP hydrolysis. Depending on the rate of nucleotide exchange, this could limit the rate of formation of ParA2-ATP dimers. We investigated nucleotide exchange kinetics by preincubating ParA2 with unlabeled AXP and rapidly mixing with MANT-AXP. The fluorescence increase indicates that pre-bound ATP or ADP cofactors on ParA2 dimers exchange with free MANT-ATP or MANT-ADP. The pseudo-first-order rate constants of nucleotide exchange, k_on_ and k_off_, were determined by titrations of increasing MANT-nucleotide concentrations (Figure 3F). ADP→ATP exchange showed a similar k_on_ (6.5×10^−4^ μM^−1^s^−1^) to ADP→ADP exchange (6.1×10^−4^ μM^−1^s^−1^), although much slower than ATP→ATP exchange (1.4×10^−3^ μM^−1^s^−1^). In comparison, k_on_ values for nucleotide exchange (Figure 3F) are much slower than for nucleotide binding (Figure 3B, D). The slow rates for nucleotide exchange correspond to ADP release and subsequent MANT-AXP binding kinetics. The observed nucleotide exchange rates k_obs_ also increased with increasing ParA2 concentrations, indicating a higher order dependence on protein concentrations (Table S1). Based on these data, the rate-limiting step in ParA2 ATPase cycle may be ADP→ATP exchange.

### ATP provides for maximum stability of ParA2 structure

As nucleotide exchange occurs at a very slow rate, we wanted to investigate if nucleotide binding induces a conformational change in ParA2. We first examined the effects of various adenine nucleotides on ParA2 secondary structure and conformational stability using circular dichroism spectroscopy. The presence of adenine nucleotides shifted the CD spectra considerably toward higher molar ellipticity (θ) at 208 nm and slightly at 220 nm, indicating a decrease in proportion of α-helices (Figure 4A). Concurrently, a slight decrease at 218 nm showed an increase of β-sheets. The overall spectra resulted in a decrease of ParA2 helicity by 10% with ATP, and a decrease by 20% with ATPγS and ADP, indicating conformational changes of ParA2 dimers upon nucleotide interactions. A similar spectral shift and loss of helicity was also observed for F SopA, suggesting analogous conformational changes upon nucleotide-binding (Libante *et al*, 2001). In contrast, P1 ParA showed an increase in helicity (4-5%) upon ADP-binding (2, 7). The effect of adenine nucleotides on ParA2 structural stability was examined by thermal melts and the molar ellipticity (θ) monitored at 220 nm between 23°C and 63°C (Figure 4B). The denaturation of ParA2 occurred at Tm = 44°C and was irreversible as ParA2 precipitated at the end of the heating cycle. ATP stabilized ParA2 structure at most by 20%, conferring a maximal T_m_ = 53°C. Both ATPγS and ADP raised T_m_ by 12% to 50°C. AMPPnP or ATP in the absence of Mg^2+^ had no effect on ParA2 stability. A similar effect of ATP and ADP on T_m_ was also observed for P1 ParA and F SopA (Davey & Funnell, 1997; Libante *et al*, 2001), indicating that this may be a common feature among chromosome and plasmid ParAs, where nucleotide-binding stabilizes ParA conformation in general although the extent of stabilization may vary depending on ParAs.

**Figure 4.**
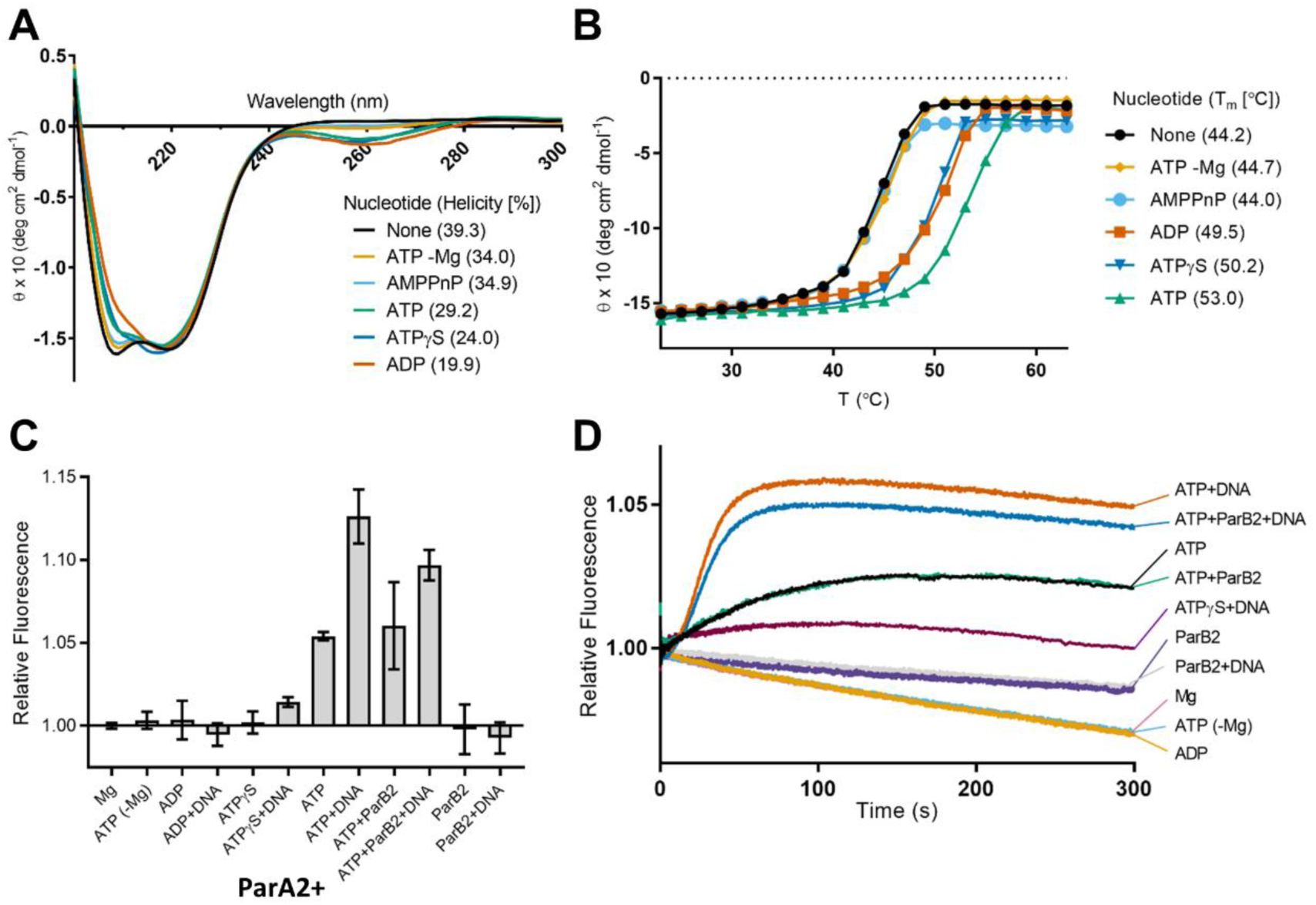
ParA2 conformational change induced by adenine nucleotides and DNA. **(A)** CD spectra of ParA2 under different nucleotide conditions. Surface helicity (%) for each condition is indicated. 5 µM ParA2 was prepared in 10 mM Tris-HCl pH 8, 5 mM MgCl2, and 1 mM adenosine nucleotide when present. **(B)** The effects of adenosine nucleotides on ParA2 stability. ParA2 changes in secondary structure monitored by CD at 220 nm (θ_220_), with an averaging time of 8 s, from 24 °C to 63°C. Samples were prepared as in **(A)**. Samples were equilibrated for 1 min prior to measurement with 2 °C increments. Relative ParA2 Tm values under different nucleotide conditions are shown in the table. **(C)** Changes in steady state ParA2 tryptophan fluorescence. ParA2 (0.6 mM) incubated in Buffer B with nucleotide (1 mM) at 23°C for 400 s before relative fluorescence change was measured. In measurements without MgCl_2_, the buffer contained 0.1 mM EDTA instead of MgCl_2_. **(D)** Kinetics of ParA2 tryptophan fluorescence change. 0.6 µM ParA2 in Buffer B at 2x final concentration in syringe 1 was rapidly mixed with 1 mM nucleotide (as indicated), 0.1 mg ml^−1^ DNA and 0.6 µM ParB2 in Buffer B at 2x final concentration at 23°C in syringe 2 of the stopped-flow apparatus.

### ParA2 undergoes a slow conformational change to ParA2*-ATP dimer

To further investigate structural changes of ParA2 upon interactions with cofactors, we took recourse to measuring intrinsic tryptophan fluorescence of the protein. ParA2 monomer has six tryptophan residues at positions 111, 224, 267, 287, 308 and 401 in the C-terminal domain. Sequence alignment with P1 ParA shows that residue W224 of ParA2 corresponds to W216 of P1 ParA, located on the α-helix 11, close to the P1 ParA-ADP dimer interface (Vecchiarelli *et al*, 2010; Dunham *et al*, 2009). We first measured the steady state fluorescence in the presence of either adenine nucleotides, DNA or ParB2, to determine if any of the cofactors could induce a fluorescence change. ParA2 tryptophan fluorescence increased by 5% in the presence of ATP and Mg^2+^ (Figure 4C, bar 7) relative to controls without ATP or Mg^2+^ (Figure 4c, bars 1, 2). ADP and ATPγS had little effect on the fluorescence change, indicating that ParA2 undergoes a conformational change that is ATP-specific (Figure 4C, bars 3, 5). The presence of DNA increased fluorescence by 13% in the presence of ATP, implying a further conformational change when ParA2 dimers associate with DNA (Figure 4C, bar 8). Although ATPγS did not initially cause a change in ParA2 tryptophan fluorescence, DNA was able to effect a slight increase, suggesting that DNA induces a ParA2 conformational change that maybe different from that with ATP (Figure 4C, bar 6). As ParB2 interacts directly with ParA2 to stimulate ATP hydrolysis, we wanted to investigate if ParB2 has an effect on ParA2 structure. ParB2 has two tryptophan residues at positions 246 and 268. However, controls with ParB2 alone showed negligible fluorescence change in the presence of ATP and DNA, demonstrating that ParB2 could be used as a cofactor in this assay without interfering with ParA2 tryptophan fluorescence (Figure 4C, bars 11 and 12). Interestingly, we found that the fluorescence change of ParA2 with ATP remained similar in the presence of ParB2. Although the fluorescence increase with DNA was slightly lowered by ParB2, suggesting a separate conformation of ParA2-DNA complex when interacting with ParB2.

In order to investigate the kinetics of ATP-induced conformational change of ParA2, we monitored the ParA2 tryptophan fluorescence over time using stopped-flow. Consistent with the steady state measurements, the controls of ParA2 without ATP or Mg^2+^ showed a prolonged decrease in fluorescence that is attributed to photobleaching (Figure 4D). Similarly, the lack of fluorescence change by ADP and ATPγS showed that they did not induce any conformational change despite binding to ParA2 dimers. On the other hand, ParA2 with ATP showed a surprisingly slow hyperbolic increase in tryptophan fluorescence to reach an apparent steady state in ∼180 s (Figure 4D). The curves fitted well to single-exponential with similar time constants τ = 57–68 s, hence the rates were not shown to be dependent on ParA2 concentrations. (Figure S4 and Table S2). The rates of fluorescence increase is much slower than MANT-ATP binding kinetics by at least 6-fold (Table S2). We hypothesize that this fluorescence change is due to a slow conformational change of ATP-bound ParA2 dimers to a distinct intermediate state, ParA2*_2_-ATP_2_. As ATP binding and nucleotide exchange have faster observed rates, we hence inferred the slow conformational change to ParA2*_2_-ATP_2_ to be the rate-limiting-step in ParA2 ATPase cycle.

### DNA modulates ParA2-ATP conformational kinetics

In the presence of DNA, ParA2 tryptophan fluorescence intensity showed an initial lag phase before increasing by more than 2-fold higher than with ATP, reaching steady state by ∼80s (Figure 4D). The lag phase was most likely due to initial nucleotide-binding, followed by the conformational change to ParA2*-ATP dimers. Hence, the curve was fitted with a single exponential with the initial lag phase excluded. Strikingly, compared to with ATP alone, the presence of DNA doubled the rate at 0.6 μM ParA2 and further sped up over 4-fold at higher ParA2 concentrations (τ=15–30 s)(Table S2). There is also a slight fluorescence increase with ATPγS, indicating that DNA induces ParA2 conformational change that is not dependent on hydrolysis. This implies that interactions with DNA activates the rate of conformational change of ParA2-ATP dimers, and lowers the energy barrier toward the ParA2*-ATP dimer intermediate state that is primed for DNA-binding. The rate and extent of fluorescence dependence on ParA2 concentrations also suggest higher order ParA2 interactions with DNA undergoing cooperative conformational transitions (Figures 4D, S4C and Table S2). Notably, *Vc* ParA2* intermediate appears to be analogous to P1 ParA* that represents the active DNA-binding state (Vecchiarelli *et al*, 2010). However, despite both ParA homologs sharing this rate-limiting step, the conformational change of *Vc* ParA2 reached steady state about 5-fold faster compared to P1 ParA. These data demonstrate that ParA2 is able to switch more quickly from a non-binding state (ParA2_2_-ATP_2_) to an active DNA-binding conformation (ParA2*_2_-ATP_2_). We infer that this is the key feature that distinguishes between plasmid and chromosomal ParAs, enabling *Vc* ParA2 to rebind DNA more quickly and to be more dynamic in exploiting and patterning the nucleoid as a scaffold for Chr2 segregation.

We next investigated the kinetics of ParA2 tryptophan fluorescence change by ParB2. Surprisingly, we found that ParB2 did not change the rates of fluorescence increase compared to with ATP, with or without DNA. Although ParB2 dampened the extent of fluorescence increase by DNA (Figure 4D, Table S2). From these data, we infer that DNA is the primary cofactor in modulating ParA2 conformational change to the DNA-binding state since ParA2-ParB2 interactions did not appear to change the conformational kinetics of ParA2*-ATP dimer formation. We hypothesize that ParB2 interacts with ParA2 by inserting an Arginine finger or α-helix at the ParA2 dimer interface to stimulate ATP-hydrolysis (Zhang & Schumacher, 2017; Volante & Alonso, 2015; Leonard *et al*, 2005). Based on the overlay of TP228 ParA-ParB and pNOB8 ParA-DNA crystal structures, the location of ParB N-terminal helix at the ParA dimer interface clashes with the DNA-binding region (Zhang & Schumacher, 2017). Thus, it was reported that ParB stabilizes ParA nucleotide sandwich dimer but not the DNA-binding state. However, when we overlaid TP228 ParA-ParB (5U1G)(Zhang & Schumacher, 2017) and Hp Soj-DNA crystal structures (6IUC)(Chu *et al*, 2019), which is more homologous to P1 ParA and *Vc* ParA2 at the DNA binding interface, we found that the ParB helices do not clash with ParA DNA-binding regions. Instead, we deduce that although ParB2 does not alter the kinetics of ParA2 conformational change, ParB2 helices interact with ParA2-DNA complexes at the dimer interface, stimulating ATPase activity and lowering the extent of conformational change.

### ParA2*-ATP dimers bind DNA cooperatively

It was shown previously that *Vc* ParA2 formed structured filaments on DNA with ATP, or ADP or in the absence of nucleotides (Hui *et al*, 2010). However, based on our data, we did not observe any ParA2 conformational change with ADP or without nucleotides, in the presence of DNA. To evaluate the nucleotide-dependence of ParA2-DNA binding activity, we performed EMSA using Cy3-labeled 62-bp double-stranded DNA and the adenine nucleotides. The agarose gels showed a sharp retarded band of ParA2-DNA complexes with increasing intensities at higher ParA2 concentrations (Figure 5A). The binding curves showed that ParA2 had similar affinities with ATP (*K_d_* = 46 nM) and ATPγS (*K_d_* = 34 nM), demonstrating that ATP hydrolysis is not required for DNA binding (Figure 5B) or conformational change (Figure 4D). The binding curves were sigmoidal, showing that ParA2 binds DNA cooperatively in the presence of ATP, with a Hill coefficient (n) = 4. This implies that two dimers of ParA2*-ATP were bound to 62-bp DNA (∼30 bp/dimer), corroborating the propensity of ParA2 to form oligomers or higher-order complexes on DNA via dimer-dimer interactions. In the presence of ADP, ParA2 could not achieve full binding and had 8-fold lower affinity for DNA (*K_d_* = 378 nM). In the absence of nucleotide, the bands were smeared throughout the lanes, showing that ParA2 by itself has a very low affinity for DNA (*K_d_* = 1 μM) and was not able to form stable complexes (Figure 5A, B). To test for ParA2 dissociation from DNA, unlabeled sonicated salmon sperm DNA (sssDNA) was added to preformed ATP-bound ParA2-DNA complexes (Figure S5). sssDNA facilitated disassembly of the complex with increasing sssDNA concentration. This was similarly observed in the presence of ATPγS, indicating that ParA2 is also able to dissociate from DNA without ATP-hydrolysis. Together, the results show that upon ATP binding, ParA2 dimer undergoes a slow conformational change to ParA2*-ATP dimer, a DNA-binding competent state; DNA catalyzes this slow step and licenses ParA2 to cooperatively bind onto DNA to form higher order [ParA2*_2_-ATP_2_]_n_ oligomers via dimer-dimer interactions.

**Figure 5.**
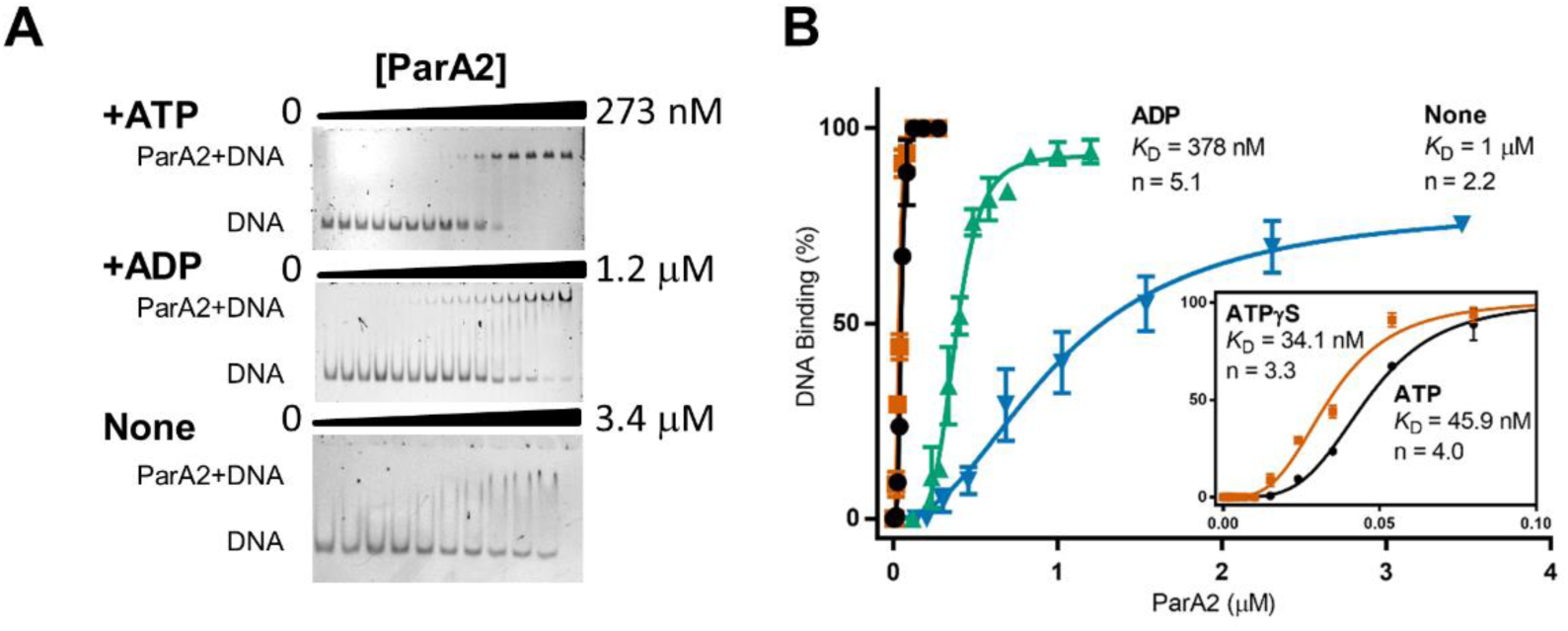
ParA2 binds DNA cooperatively with ATP. EMSA of ParA2 binding to Cy3-62-bp DNA (5 nM). EMSA of ParA2 at increasing concentrations as indicated were titrated with 5 nM Cy3-labeled 62-bp DNA in Buffer A, in the presence of 2 mM ATP (top), ADP (centre) or without nucleotides (bottom). The samples were run in a 5% PAGE gel in TBM buffer. The positions of free DNA and the ParA2-DNA complexes are indicated on the left. **(B)** ParA2-DNA binding affinity. % DNA binding was calculated using Image J. Data was plotted, with indicated *K*_D_s and Hill coefficients (n) under each condition. **(A)** ParA2 at increasing concentrations as indicated were titrated with 5 nM Cy3-labeled 62-bp DNA in Buffer A, in the absence of nucleotides. The samples were run in a 5% PAGE gel in TBM buffer. The positions of free DNA and the ParA2-DNA complexes are indicated on the left.

### ATP stimulates ParA2*-GFP binding and dissociation on DNA carpet

We wanted to directly observe ParA2-DNA interactions and its dependence on nucleotides to investigate its kinetic properties. We coated a two-inlet flow cell surface with a DNA carpet acting as a biomimetic chromosome and visualized with a TIRF microscope ParA2 fused at its C-terminal to GFP. The two-inlet flow cell allows us to instantly switch between flowing sample solution and wash buffer to monitor real-time binding and dissociation kinetics of ParA2-GFP on the DNA carpet. DNA-binding and dissociation activities of ParA2-GFP were checked to be similar to wild-type ParA2 (Figure S5). We preincubated 10 μM ParA2-GFP with various adenine nucleotides and diluted the mixture 10-fold (to 1 μM ParA2-GFP) before infusing into the flow cell. This helps to saturate nucleotide-binding to ParA2 dimers so that the observed kinetic changes could be attributed primarily to DNA-binding. In the presence of ATP, ParA2-GFP bound to the DNA carpet exponentially and instantly, with a time constant τ = 19 s and the binding reached steady state within 1 min (Figure 6A, Table S3). When we switched to flow wash buffer at 6 min, ParA2-GFP intensity dropped almost immediately, dissociating from the DNA carpet at a rate twice faster than binding. At the concentration used, ParA2-GFP coated the DNA carpet thoroughly and uniformly, and dissociated evenly with wash buffer, showing highly efficient reversible binding (Movie S2). In the presence of ATPγS, although ParA2-GFP bound the DNA carpet to a similar extent as with ATP, the observed rates of binding and dissociation were slower (Table S3). This supports our EMSA data that ATP hydrolysis is not required for ParA2-DNA binding and dissociation but stimulates rates of DNA association (3-fold) and dissociation (8-fold). When infused with ADP or without nucleotides, ParA2-GFP showed negligible binding to DNA carpet, contrary to previous EM studies (Hui *et al*, 2010). Although we saw DNA binding with EMSA, this was at low affinity. We believe that in the flow cell, due to the presence of high concentrations of nonspecific DNA relative to protein concentrations, this condition mimics intracellular environment more closely.

**Figure 6.**
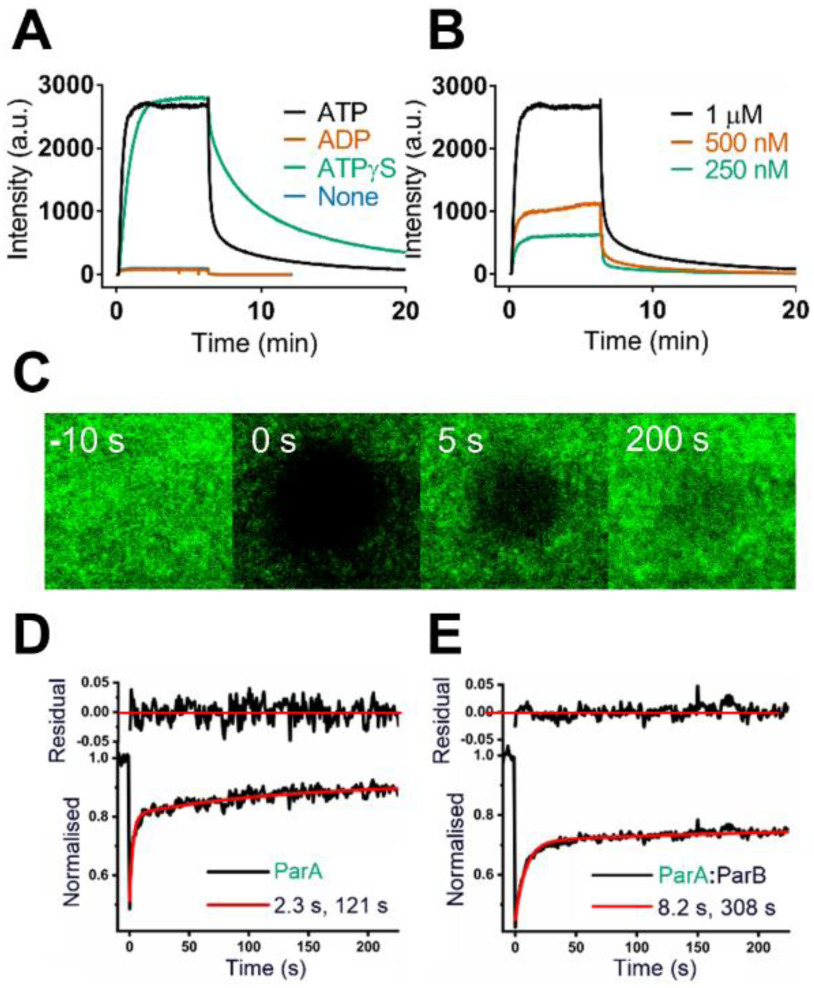
Fast exchange of ParA2 on DNA carpet stabilized by ParB2. **(A)** Binding and dissociation curves of ParA2 on DNA carpet. ParA2-GFP (10 μM) was preincubated in the presence or absence of ATP, ADP or ATPγS (2 mM). Protein was diluted to 1 μM before infusion into DNA-carpeted flowcell until steady state and switched to wash buffer at 6 min. **(B)** ParA2 binds DNA carpet cooperatively at higher protein concentrations. ParA2-GFP (10 µM) was preincubated with ATP (1 mM) prior to dilution to final concentrations shown. Sample was infused into flowcell and washed as in (A). All binding and dissociation rates are listed in Table S3. **(C)** Time-lapse images of ParA2-GFP (low density) recovery on DNA carpet after photobleaching at 0 s. **(D)** Representative FRAP curve (black line) of ParA2-GFP was fitted to a double-exponential (red line). The fitted time constants correspond to a fast species (2.3 s) and slow species (121 s). **(E)** Representative FRAP curve (black line) of ParA2-GFP with ParB2 at A:B 1:2 ratio was fitted to a double exponential (red line). The fitted time constants correspond to a fast species (8.2 s) and slow species (308 s). All FRAP data are listed in Table S4.

We next studied how varying protein concentrations affect ParA2-DNA binding. 10 µM of ParA2-GFP was preincubated with 2 mM ATP and diluted to its final protein concentrations of 250 nM-1 µM before infusion (Figure 6B). Intensities of ParA2-GFP at steady state increased proportionately from 250 to 500 nM but showed higher order ParA2-GFP concentration dependence at 1 µM, indicative of a cooperative binding process. All three concentrations had a similar binding rate of 0.05 s^−1^ (3 min^−1^) and dissociation rates of 0.11–0.19 s^−1^ (6.25–8.6 min^−1^) (Table S3). We found that the dissociation rates are 5–8-fold faster than plasmid ParAs, P1 ParA (0.8–1.8 min^−1^) and F Sop A (1.9 min^−1^) (Vecchiarelli *et al*, 2010; Hwang *et al*, 2013). The dissociation rates were also sped up by the presence of ParB2 and DNA in the wash buffers, indicating intersegmental transfer of ParA2-GFP between DNA and stimulated hydrolysis and dissociation by these cofactors (Figure S6).

From our *in vivo* images (Figure 1 and Movie S1), *V. cholerae* cells take less than 30 s for ParA2-GFP to dissociate from the nucleoid and transition across to the opposite cell pole and rebind there. The rapid dissociation rates measured here corroborate with the fast dynamic oscillations of ParA2-GFP observed in the cell. In contrast, when we mixed ParA2-GFP and ATP just before infusing into the flow cell in an ‘ATP-start’ experiment, there was an initial lag time of 1 min before intensity increased (Figure S7), which we attribute to ATP-binding. Prominently, the extent of binding dropped 5–10-fold compared to pre-incubation experiments. ParA2-GFP binding rates were also slower by up to 3-fold than preincubation experiments. These results support our tryptophan fluorescence data, showing a time delay switch upon ATP-binding for ParA2-GFP to transition to an active DNA-binding conformation.

### ParA2*-GFP rapid exchange on DNA carpet is stabilized by ParB2

Fogel and Waldor proposed that *V. cholerae* Chromosome 1 segregates via a mitotic-like mechanism where ParA1 polymerizes to form a dynamic filament to pull one of the replicated ParB1 bound centromeric loci toward the new cell pole (Fogel & Waldor, 2006). Hui et al. have shown using negative stains that *Vc* Chr2 ParA2 forms stable filaments on DNA with and without nucleotides (Hui *et al*, 2010). However, these are inconsistent with the fast rates measured by our DNA carpet binding experiments. We wanted to verify if Chr2 ParA2 is able to form stable filaments on DNA. To this end, we used fluorescence recovery after photobleaching (FRAP) to probe the exchange of ParA2-GFP on the DNA-carpet. ParA2-GFP was infused into the flow cell to two different densities (28% and 100% at steady state), which are in excess of estimated physiological density at 1% density (Hwang *et al*, 2013). A spot was photobleached <1 s on the ParA2-GFP-coated carpet and the fluorescence recovery monitored (Figure 6C). If ParA2-GFP forms stable filaments on the DNA carpet, we would expect a large fraction of unrecovered or slowly recovering species. However, at 28% density, we found only a small fraction of immobile species (13%). Majority of ParA2-GFP recovered very rapidly with a fast species of τ = 2.3 s (64% at 0.43 s^−1^) and a smaller fraction of slower species of τ = 121 s (23% at 0.008 s^−1^) (Figure 6D and Table S4). As most of the proteins were DNA-bound and recovered by exchanging with ParA2-GFP from the solution phase, the photobleaching recovery rate depends on the rate of protein unbinding. At 100% density or steady state binding, ParA2-GFP recovery time increased to τ = 8.5 s (44% at 0.12 s^−1^) and the fraction of fast species shifted towards slower and immobile species. This suggests that with increasing protein/DNA ratio, there is an increasing population of ParA2-GFP that is forming higher order complexes on DNA that is exchanging slower on the carpet. The exchange rate is also consistent with the dissociation rate constants determined from the wash experiments above (Table S2). When compared to plasmid ParAs, the rate of recovery of the major fraction of *Vc* ParA2 is 15-fold and 6-fold faster than P1 ParA (1.8 min^−1^) and F Sop A (4.7 min^−1^), respectively (Hwang *et al*, 2013; Vecchiarelli *et al*, 2013). These data highlight the faster exchange rate of ParA2-GFP on DNA compared to plasmid ParAs.

As ParB2 stimulates ParA2 ATPase activity, we wanted to test if ParA2 exchange on DNA is also affected. We found that ParB2 lowers ParA2-GFP density on DNA carpet and inhibits protein exchange, increasing the recovery time constants by up to 4-fold at higher ParB2:ParA2 ratio (Table S4). In addition, the dominant fraction of faster species was reduced in the presence of ParB2, converting to the immobile fraction. This indicate that ParB2 stabilizes ParA2 binding on DNA and slows down its dissociation. A similar effect was also reported for P1 ParA and F SopA (Hwang *et al*, 2013; Vecchiarelli *et al*, 2013), implying two separate populations of ParA: one that is DNA-bound and another that is interacting with ParB on the partition complex. We infer that ParB2, when bound to the partition complex at high concentrations, slows down ParA2 exchange on the partition complex and its vicinity, allowing for longer-lived depletion of ParA2 from DNA. Collectively, our data – fast ParA2-GFP recovery rates, fast DNA dissociation rates and the lack of observed stable filaments on DNA carpet – do not support the model that ParA2 forms extended filaments on DNA that is stable enough to exert a pulling force on the chromosome. Instead, the data support a diffusion-ratchet model where ParA2 cooperatively binds Chr2 as dynamic oligomers to pattern the nucleoid, mediating its movement and bidirectional segregation.

## DISCUSSION

### Kinetic pathway of ParA2 ATPase cycle and DNA binding

We have characterized for the first time, the kinetic pathway of the ATPase cycle and DNA binding of a chromosomal ParA that is involved in segregating Chr2 of *V. cholerae* (Figure 7). Chr2 ParA2 forms dimers at low concentrations spontaneously. Upon binding ATP (*k_1_*, *k_−1_*), the ParA2_2_-ATP_2_ undergoes a slow conformational change from a closed sandwich dimer to ParA2_2_*-ATP_2_ state (*k_2_*), an open dimer that licenses the dimers to bind DNA. This slow transition is accelerated in the presence of DNA by 2 to 5-fold. Once competent for DNA-binding, ParA2_2_*-ATP_2_ loads onto DNA and induces cooperative binding of more ParA2 dimers via dimer-dimer interactions to form higher-order oligomers on DNA (*k_3_*, *k_−3_*). Although, ParB2 did not appear to influence the rate of ParA2 conformational change, ParB2 inhibited ParA2 exchange on DNA as it stabilizes ParA2-DNA interactions. Upon ATP hydrolysis and two phosphate release, ParA2_2_-ADP_2_ dissociates from DNA (*k_4_*). The presence of competitive DNA and ParB2 activates the rate of DNA dissociation, due to stimulated hydrolysis. Once ADP dissociates from ParA2 dimers (*k_5_*, *k_−5_*), the cycle restarts and ParA2 dimers diffuse away and are ready to rebind available ATP for the next round of DNA-binding. ParA2 dimers can undergo nucleotide exchange (*k_6_, k_−6_*) at much slower rates than ATP-binding. Although the rate of ParA2 recruitment on DNA is primarily determined by the rate-limiting conformational change to its active state, there is also a secondary dependence on the nucleotide exchange rate. Particularly, once DNA activates ParA2 conformational change, the slow nucleotide exchange step controls ParA2 recruitment and dynamic localization. Another oscillation system, MinCDE proteins have been previously modeled to show that nucleotide exchange rates control the spatial distribution of MinD rebinding and oscillations (Halatek & Frey, 2012). Hence, we believe that the ParA2 dynamic patterns on the nucleoid are spatiotemporally regulated by the rates of these two processes.

**Figure 7.**
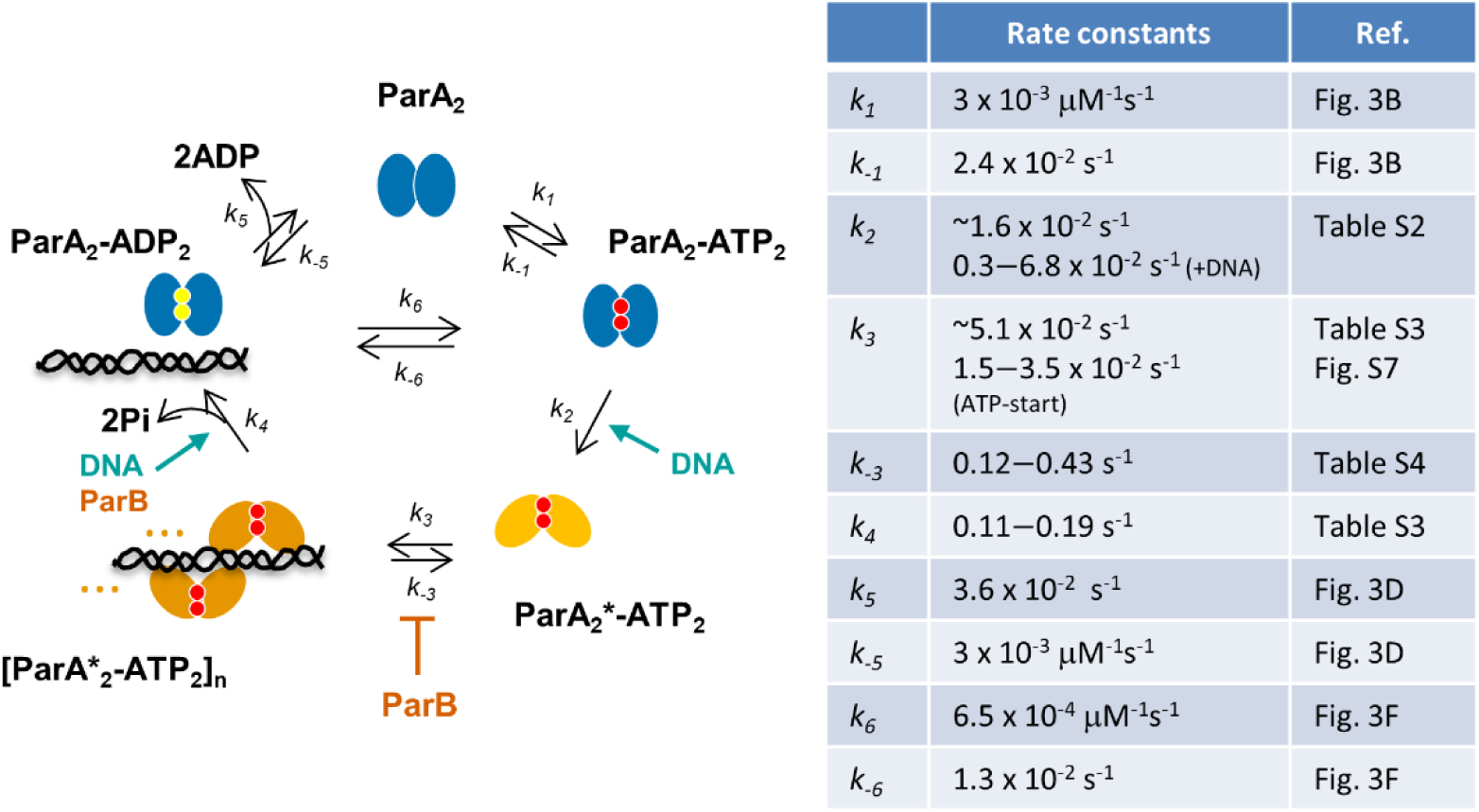
Kinetic pathway and table of rates for ATP control of ParA2-DNA interactions. ParA2 dimers bind ATP to form a closed sandwich dimer (*k_1_*, *k_−1_*). ParA2_2_-ATP_2_ undergoes a slow conformational change to active ParA2_2_*-ATP_2_ state (*k_2_*), an open dimer that licenses ParA2 dimers to bind DNA. This slow transition is accelerated in the presence of DNA. Once competent for DNA-binding, ParA2_2_*-ATP_2_ loads onto DNA and induces cooperative binding of more ParA2* dimers via dimer-dimer interactions to form oligomers on DNA (*k_3_*, *k_−3_*). ParB2 inhibits ParA2* exchange on DNA as it stabilizes ParA2*-DNA interactions. ParA2_2_-ADP_2_ dissociates from DNA (*k_4_*) upon ATP hydrolysis that is stimulated by DNA and ParB2. Once ADP dissociates from ParA2 dimers (*k_5_*, *k_−5_*), the cycle restarts and ParA2 dimers diffuse away and are ready to rebind ATP for the next round of DNA-binding. ParA2 dimers can undergo nucleotide exchange (*k_6_, k_−6_*) at much slower rates than ATP-binding. ParA2 recruitment on DNA and its dynamic oscillations are primarily controlled by the rate-limiting conformational change to its active state, and the secondary control by the nucleotide exchange rate. (*k_2_* and *k_3_* are the averaged rates from different protein concentrations.)

### ParA2 gradient formation on nucleoid DNA

Based on our biochemical characterization, we propose a model for ParA2 gradient formation on *V. cholerae* nucleoid and how this could be involved in transporting Chr2 from the cell centre to quarter positions of the cell before cell division. The rate of ParA2 rebinding to DNA is dependent on three factors: the slow conformational change of ParA2 to the active ParA2* state that is stimulated by DNA, the nucleotide exchange rate that is dependent on higher order ParA2 concentrations and slowing down of DNA exchange rate by ParB2. Overall, the rate of ParA2 rebinding on DNA is directly correlated to the concentration of Par proteins or the concentration ratios of ParA2:ParB2 in the cell. In a young cell when *ori2* is at midcell, the concentration of ParB2 is highest on the *par* locus at the midcell and diminishes towards the cell poles. From our *in vivo* images, ParA2 concentrations are highest either towards one or the other pole. Hence, over time average, ParA2 concentration will be highest at the cell poles and lowest at midcell. We hypothesize that upon ParB2-stimulated ParA2 release from DNA, the rate of ParA2 rebinding on the nucleoid will be slowest at the midcell where the highest concentration of ParB2 is localized on *parS2*, and fastest toward the cell poles where concentration of ParB2 is the lowest. This distribution of ParA2 rebinding rates induced by ParB and DNA concentration ratios set up a spatiotemporal gradient of ParA2 concentration across the cell. For F and TP228 plasmids, their motion coordinate with ParA migration by chasing the receding ParA gradient. This is primarily based on the diffusion-ratchet mechanism, where the plasmids ratchet along a moving ParA gradient using collective dynamic ParA-ParB interactions (Hwang *et al*, 2013; Vecchiarelli *et al*, 2013, 2014). We predict Chr2 uses a fundamentally similar mechanism for segregation. However, because ParB2-*parS2* locus remains largely stationary relative to the dynamic oscillating ParA2 wave. We speculate that in young cells, ParA2 wave cooperatively binds onto the nucleoid as dynamic oligomers and constantly sweeps from pole-to-pole to ‘tug’ at ParB2-*parS2* locus. This fine-tunes and maintains the locus at midcell position with each periodic oscillation. Upon replication, the doubling of ParB-*parS2* loci activates further depletion of ParA2 at midcell, redistributing the ParA2 wave towards elongating cell poles. This then drives the loci to bidirectionally segregate toward quarter cell positions in older predivisional cells. Once the division septum is formed, ParA2 oscillation reestablishes between the new cell poles in the daughter cells and repositions the *par* loci back at midcell.

### Comparison of chromosomal and plasmid ParAs

Our studies of *Vc* ParA2 revealed significant differences in the properties between chromosomal and plasmid ParAs. Compared to P1 ParA and other plasmid ParAs, ParA2 forms dimers spontaneously unlike the ATP-dependent dimerization of plasmid ParAs. ParA2 has a faster ATP binding rate and higher specific ATPase activity; the relative stimulatory effects by ParB2 and DNA are also more pronounced (Leonard *et al*, 2005; Barillà *et al*, 2005; Lim *et al*, 2014; Lee *et al*, 2006; Scholefield *et al*, 2011; Davis *et al*, 1992). Despite *Vc* ParA2 and P1 ParA having the same rate limiting step, ParA2 undergoes a faster conformational change to the active state, leading to faster DNA-rebinding. We infer that in the cell, the fast exchange rates of ParA2 on the nucleoid will lead to fast on and off rates with its cognate partner ParB2. These faster transient interactions between ParA2 and the nucleoid, as well as ParA2 and ParB2-*parS2*, allow for the chromosome partition complex to be more dynamic in interacting with the nucleoid, relative to plasmids. Overall, the faster reaction rates imply that *Vc* ParA2 is a more efficient and robust enzyme, and selection of its features may have been necessitated to efficiently segregate a larger cargo of Chr2 in coordination with the cell cycle. We believe that the kinetic features of ParA2 could be shared among the chromosomal ParAs and it would be important to test that.

## MATERIALS AND METHODS

### Buffers

**Buffer A**: 50 mM Tris-HCl pH 7.5, 100 mM NaCl, 10 mM MgCl_2_, 10% glycerol, 100 mg/ml BSA, and 1 mM DTT. **Buffer B**: 50 mM Tris-HCl pH 7.5, 150 mM NaCl, 5 mM MgCl_2_. **Par Buffer**: 50 mM Tris (pH 7.5), 100 mM NaCl, 5 mM MgCl_2_, 10% (v/v) glycerol, 1 mM DTT, 0.1 mg/ml α-casein.

### Construction of plasmid encoding ParA2-GFP fusion

See supplementary information.

### Microscopy of ParA2 in *V. cholerae*

Cells were grown in 1X M63 medium supplemented with 1 mM CaCl_2_, 1 mM MgSO_4_, 0.001% vitamin B1, 0.2% fructose, 0.1% casamino acids at 30°C until an OD at 600 nm of 0.3. When required, kanamycin was added to a final concentration of 12.5 µg/ml. Expression of ParA:GFP was induced by adding 0.0008% arabinose for 1 hour at 30 C with shaking. 10 µl of the culture was plated on the center of a glass P35 dish (MatTek corporation, Ashland, MA), and overlaid with a 1% agarose disc prepared with the same M63 medium described above but supplemented with 0.02% arabinose. Images were taken every 30 secs on a Nikon Ti-Eclipse inverted microscope with Nikon 100x/1.4 Oil Plan Apo Ph3 DM lens, imageEM EMCCD camera (Hamamatsu, Japan) and Lumencor sola light engine (Beaverton, OR) set to 5% 475 nm laser output and 300 ms exposure.

### Protein expression and purification

See supplementary information.

### SEC-MALS

Samples of 40 μM ParA2 were incubated alone, or in the presence of 1.0 mM ATP or ADP, in 50 mM Tris-HCl pH 7.5, (210 mM NaCl, 5.0 mM MgCl_2_, 0.1 mM EDTA, 1.0 mM DTT, 1.0 mM NaN_3_), for 20 minutes at 37°C. SEC-MALS of ParA2 was performed with 15 μl injections into a GE Superdex 200 10/300 GL SEC column at 0.75 ml/min equilibrated and run in 50 mM Tris-HCl pH 7.5 buffer (100 mM NaCl, 5.0 mM MgCl_2_, 0.1 mM EDTA, 1.0 mM DTT, 1.0 mM NaN_3_) using a Postnova AF2000 system with PN5300 autosampler. Protein elution was monitored with a Shimadzu Prominence SPD-20AV (PN3212) UV absorbance detector, PN3621b MALS detector and PN3150 Refractive Index Detector. Data analysis was conducted with NovaFFF AF2000 2.1.0.1 (Postnova Analytics, UK Ltd.) software and values plotted in Graphpad Prism 8.0.2. For protein concentration determination, a UV_280nm_ molar extinction coefficient of 1.03 M^−1^ cm^−1^ was used and absolute molecular weights were calculated using Zimm fits. Data was averaged in triplicate.

### EMSA

A standard reaction mixture (20 µl) was prepared in Buffer A with 5 nM Cyanine 3-labeled 62-bp DNA and 2 mM of ATP, ADP, ATPγS or no nucleotide, with increasing concentrations of ParA2 as indicated. The reactions were assembled on ice, incubated for 30 min at 30°C, and analyzed by gel electrophoresis in 5% polyacrylamide gels in TBM (90 mM Tris, 150 mM Borate, 10 mM MgCl_2_). Gel electrophoresis was pre-run at 120 V for 30 min, at 4°C, in a Mini-PROTEAN Tetra Cell, and then run at 120 V for 1 h, at 4°C. Gels were imaged using a Bio-Rad ChemiDoc™ MP Imaging System using the Cy3 channel with 2 minutes exposure. Images were analyzed with ImageJ (NIH).

### Circular dichroism spectroscopy (CD)

CD experiments were performed in 10 mM Tris pH 8.0, 5 mM MgCl_2_, which was filtered and degassed to prevent oxidation in the absence of DTT. Reaction mixtures were prepared by adding 5 µM ParA2 with 2 mM ATP, ADP, AMPPNP, ATPγS, or without nucleotide, to a final volume of 230 µl. An additional sample of ATP in the absence of MgCl2 was prepared as above but in 10 mM Tris pH 8.0, 2 mM EDTA. Each sample was filtered by centrifugation using a 0.2 µm Generon Proteus Clarification Mini Spin Column (GENMSF-500), at 14,000 g for 2 min. The remaining 210 µl was incubated at 23°C for 15 min. Spectra were measured using a Jasco J-810 Spectropolarimeter in a 1 mm Hellma Analytics QS High Precision Cell. Measurements were collected from 300 to 200 nm+/− 2.5 nm, in 1 nm intervals with an 8 s integration time. The spectrum of a buffer blank with or without 2 mM nucleotide was subtracted from the ParA2 spectrum with or without the corresponding nucleotide. Each experiment is repeated at least twice and each spectra recorded is an average of 3 scans. Each experiment is repeated at least twice and each spectra recorded is an average of 3 scans. ParA2 secondary conformation was monitored by CD at 220 nm+/-2.5 nm with 8 s integration time, from 23°C to 63°C. The temperature was increased in 2°C increments, and the sample was equilibrated to each temperature for 1 min before measurement of the signal.

### ATPase activity

For pre-steady state ATPase activity measurements, 1.5 µM ParA2, 100 µM ATP and 64 nM [α-^32^P]ATP were incubated in Buffer A. Where indicated, 1.5 µM ParB2 and/or 0.1 mg/ml sonicated salmon sperm DNA were added. 10 µl reactions were assembled on ice, incubated for the indicated time periods at 37°C and quenched by the addition of 10 µl 1% SDS, 20 mM EDTA. For steady state activity assays, indicated concentrations of ParA2 were incubated in reactions set up as described above, at 37°C for 30 min. 1 µl from each sample was spotted onto a POLYGRAM CEL 300 PEI-TLC plate (Macherey-Nagel), and developed with 0.5 M LiCl (Sigma). 1 M formic acid (Alfa Aeser). Dried plates were exposed to a storage Phosphor screen and scanned with a phosphoimager (Typhoon FLA7000 IP) for quantification using ImageJ (NIH).

### Nucleotide binding, dissociation and exchange assays

Stopped flow measurements with MANT (*N*-methylanthraniloyl)-labeled nucleotides (Jena) were performed at 23°C using Stopped flow measurements with MANT (*N*-methylanthraniloyl)-labeled nucleotides (Jena) were performed at 23°C using an Applied Photophysics SX20 system. system. The excitation monochromator wavelength was set to 356 nm±1.2 nm. The emission filter on the PMT was BLP01-405R-25 (Semrock). Nucleotide binding, dissociation, and exchange experiments were performed in Buffer B with samples prepared on ice. For nucleotide binding assays, 0.6, 1.25, 2.5 μM ParA2 was rapidly mixed with 25 μM MANT-AXP and fluorescence increase was monitored over time. For pseudo-first order reaction, 0.3125, 6.25, 1.25, 2.5 µM ParA2 was rapidly mixed with 3.125, 6.25, 12.5, 25 μM MANT-AXP in buffer B and their fluorescence increase monitored. The observed binding curves were fitted with single exponential increase to determine observed rate of binding, k_obs_. Plots of k_obs_ vs. substrate concentration yielded k_on_ and k_off_ from the slopes and y-intercepts, respectively (Hulme & Trevethick, 2010). For nucleotide dissociation assay, 2.5 μM ParA2 and 5 μM MANT-AXP were pre-incubated at 23°C for 3 min, then rapidly mixed with 1 mM unlabeled AXP and their fluorescence decrease monitored. For nucleotide exchange assay, 0.625, 1.25, 2.5 μM ParA2 was preincubated at a 1:5 ratio with 3.125, 6.25, 12.5 μM unlabeled AXP, respectively, then rapidly mixed with 15.625, 31.25, 62.5 μM MANT-AXP at 5x higher concentrations than AXP. All data were averages of at least two experiments. Values were reported as relative fluorescence increase or decrease.

### Tryptophan fluorescence experiment

For steady state reactions, 0.6 µM ParA2 with 1 mM ATP, ADP or ATPγS were incubated at 23°C for 15 min in Buffer B. In the absence of MgCl_2_, a separate buffer was prepared with 0.1 mM EDTA and without MgCl_2_. Tryptophan fluorescence signal was acquired using a SpectraACQ spectrafluorimeter set at 356 nm±1.2 nm in a Hellma Analytics High Precision Cell. FluorEssence V3.5 software was used for plotting data and GraphPad Prism for data analysis. Stopped-flow measurements were performed at 23°C using an Applied Photophysics SX20 system. The excitation monochromator was set to 295 nm. The emission filter on the PMT was BLP01-325R-25 (Semrock). For kinetics experiment, 1.2 µM ParA2 was rapidly mixed with 2 mM MANT-AXP in buffer B and when present, 0.2 mg/ml DNA and 1.2 µM ParB2. Final concentrations after mixing are half of initial concentrations. All results are averages of at least two experiments. Values were reported as relative fluorescence increase or decrease.

### Flowcell coating with DNA carpet

Single-inlet and dual-inlet flowcells (with Y-channels) were coated with biotinylated lipid bilayer, neutravidin and DNA carpet as previously described (Hwang *et al*, 2013; Vecchiarelli *et al*, 2013). The following day, DNA-carpet was washed with Par Buffer and incubated for 1 h to passivate the surface with caesin. The DNA carpeted flowcell was subsequently washed with Par Buffer before sample flow.

### TIRF microscopy

A home-built prism-based TIRF microscope coupled to microfluidics was used for imaging the DNA carpet. See supplementary information for further details.

### ParA2-GFP binding and dissociation on DNA carpet

ParA2-GFP (10 µM) was preincubated in Par Buffer with 1 mM ATP, ADP or ATPγS for 30 min at 25°C. The sample was diluted to final protein concentrations as indicated with Par buffer. The sample was loaded into a 1 ml syringe (BD) and attached to inlet 1 of a dual-inlet flowcell with a Y-shaped channel. A separate syringe containing wash buffer (Par Buffer) was attached to inlet 2. The TIRF illumination field and microscope objectives were aligned to the Y-channel junction at the point of flow convergence. For ParA-GFP binding, sample and wash buffers were infused simultaneously into the flowcell at 20 µl/min and 1 µl/min respectively. After 380 seconds, sample and wash buffer flow rates were switched to 1 µl/min and 20 µl/min respectively, for protein dissociation. Data was analyzed using GraphPad Prism 7. ParA2-GFP fluorescence intensities were subtracted from background. Binding curves were fitted with ‘one phase association’ model whilst dissociation curves were fitted with a ‘two phase decay’ model.

## ACKNOWLEDGEMENTS

SC’s and ABC’s PhD studentships are funded by EPSRC Chemical Biology and BBSRC White Rose DTP (BB/M011151/1), respectively. We acknowledge Postnova Analytics UK Ltd. for the kind loan of the PN3621b MALS detector. We are grateful to Svetlana Sedelnikova for her assistance in protein purification, Rosemary Staniford for the use of her CD spectrometer, Neil Hunter lab for the use of their spectrophotometer and Jian Liu for helpful discussions. We acknowledge funding from Royal Society (RG150776), Society for Applied Microbiology and Biochemical Society. This work was supported (in part) by the Intramural Research Program of the NIH, NCI.

## AUTHOR CONTRIBUTIONS

LCH and DC designed and supervised the study. SC, RR and LCH made the constructs and strains. SC expressed and purified proteins. ABC and LCH set up the TIRF microscope. SC, ABC, PD, RR, LCH performed experiments and analyzed data. SC, ABC, RR, LCH and DC wrote the paper.

## SUPPLEMENTARY INFORMATION

**Movie 1. Binding and subsequent dissociation of ParA2-GFP on DNA carpet.**

Final concentration of 1 μM ParA2-GFP preincubated with 1 mM ATP was infused into the DNA-carpeted flowcell and imaged with TIRFM. Intensity increase shows ParA2-GFP binding DNA carpet. Flow was switched to wash buffer at 26 s (in movie). Intensity decrease shows ParA2-GFP dissociation. Data acquisition rate was 1 fps at 100 ms exposure time. Movie playback is 15 fps.

**Movie 2. FRAP of ParA2-GFP at high density bound to DNA carpet**

Final concentration of 1 μM ParA2-GFP preincubated with 1 mM ATP was infused into the DNA-carpeted flowcell until steady state (100% density). The ParA2-GFP-coated DNA carpet was photobleached at 1 s (in movie) and fluorescence recovery monitored with TIRFM. Data acquisition rate was 1 fps at 100 ms exposure time. Movie playback is 10 fps.

**Movie 3. FRAP of ParA2-GFP at low density bound to DNA carpet**

Final concentration of 1 μM ParA2-GFP preincubated with 1 mM ATP was infused into the DNA-carpeted flowcell for a shorter time before reaching steady state (28% density). The ParA2-GFP-coated DNA carpet was photobleached at 1 s (in movie) and fluorescence recovery monitored with TIRFM. Data acquisition rate was 1 fps at 100 ms exposure time. Movie playback is 10 fps.

**Figure S1.**
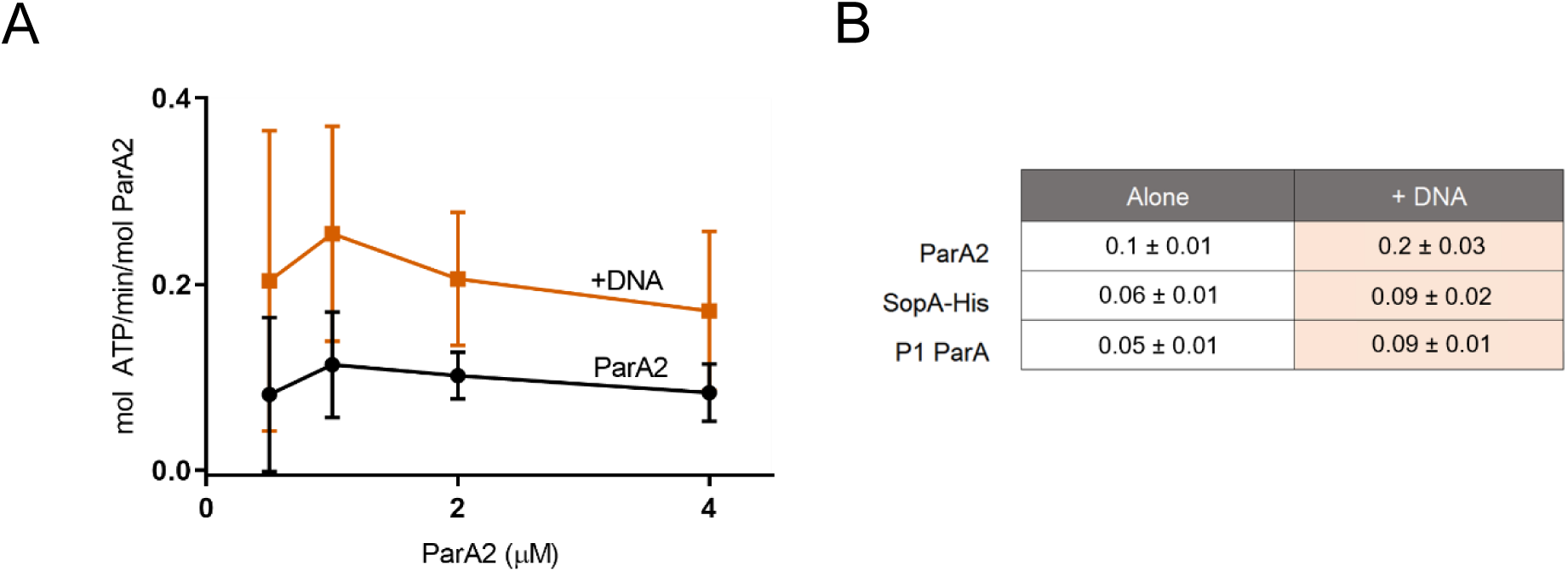
Specific ATPase activity of ParA2. **(A)** Specific ATPase activity of ParA2 with and without DNA (0.1 mg/ml) (this study). **(B)** Relative specific ATPase activity of ParA2 compared to F SopA and P1 ParA with and without DNA (MacCready *et al*, 2018). ParA2 shows higher specific ATPase activity and is more stimulated by the presence of DNA.

**Figure S2.**
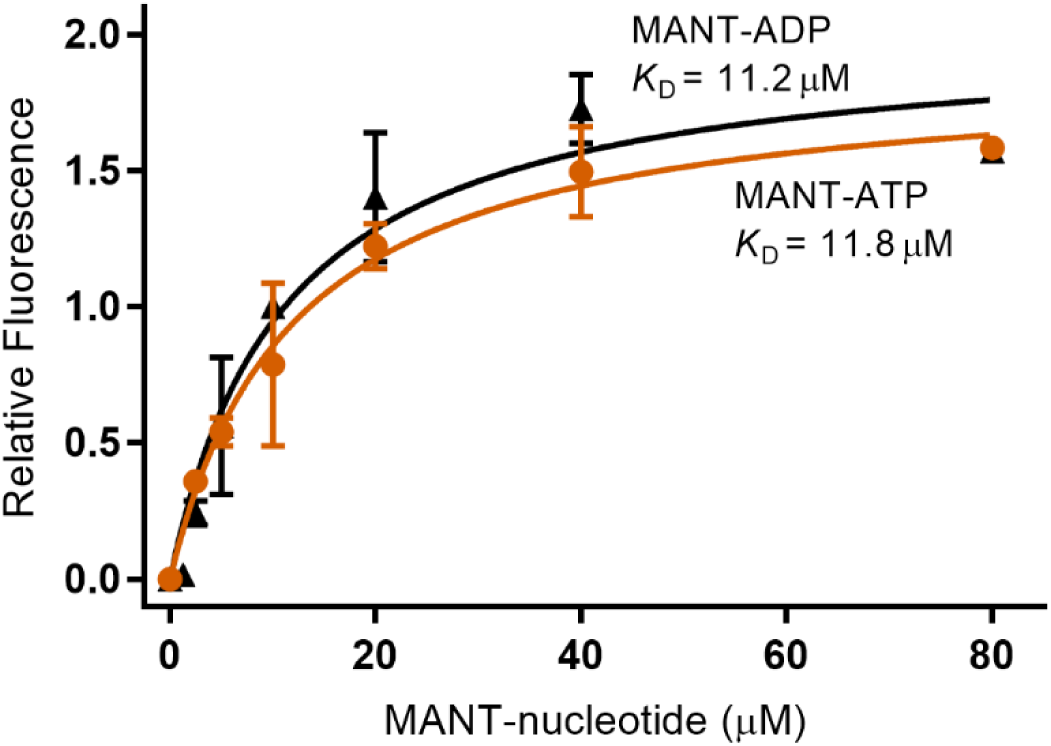
Binding curves of ParA2 and MANT-AXP. 1.5 μM ParA2 was prepared with indicated concentrations of MANT-ATP or MANT-ADP in Buffer B on ice. An initial fluorescence measurement was taken for each sample before incubating at 37°C for 20 min. The steady state fluorescence was then measured for each sample. Readings were acquired using a Fluorolog^®^-3 spectrofluorometer (Horiba Scientific) and a ‘SpectraACQ’ controller set at 356 nm ± 1.2 nm, in a ‘HellmaAnalytics High Precision Cell’. Experiments were repeated two times. The relative fluorescence change was fitted in GraphPad Prism 8, with a saturation, one-site specific binding equation, to derive *K*_D_.

**Figure S4.**
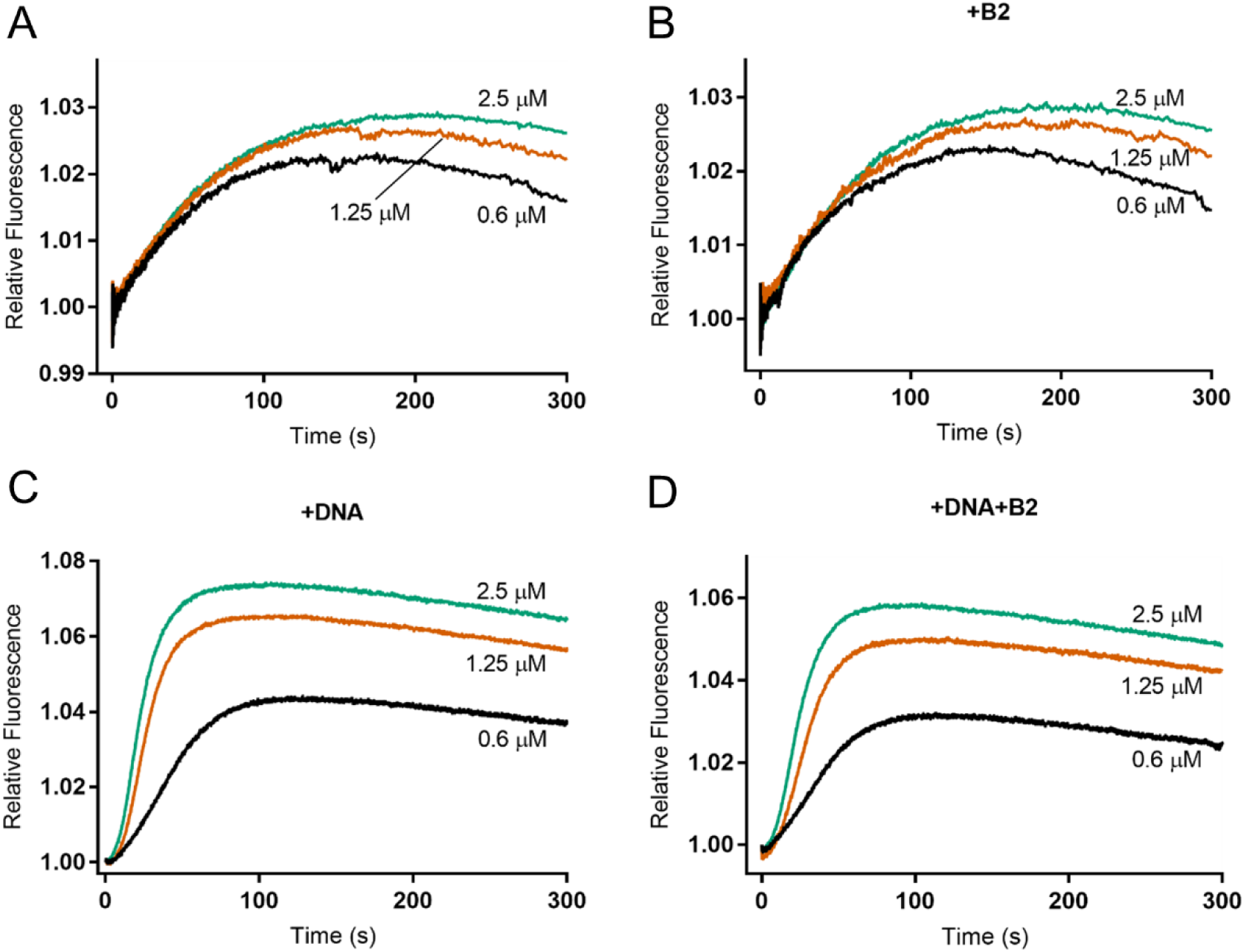
Stopped flow kinetics of ParA2 tryptophan fluorescence. **(A)** Premix 1 with indicated concentrations of ParA2 was rapidly mixed with premix 2 containing 1 mM ATP, both in Buffer B. Relative fluorescence increase was monitored over time. **(B)** ParA2 tryptophan fluorescence kinetics in the presence of ParB2. As in (A), except with 0.6 μM ParB2 added to premix 2. **(C)** ParA2 tryptophan fluorescence kinetics in the presence of DNA. As in (A), except with 0.1 mg/ml DNA added to premix 2. **(D)** ParA2 tryptophan fluorescence change in the presence of ParB2 and DNA. As in (A), except with 0.6 μM ParB2 and 0.1 mg/ml DNA added to premix 2.

**Figure S5.**
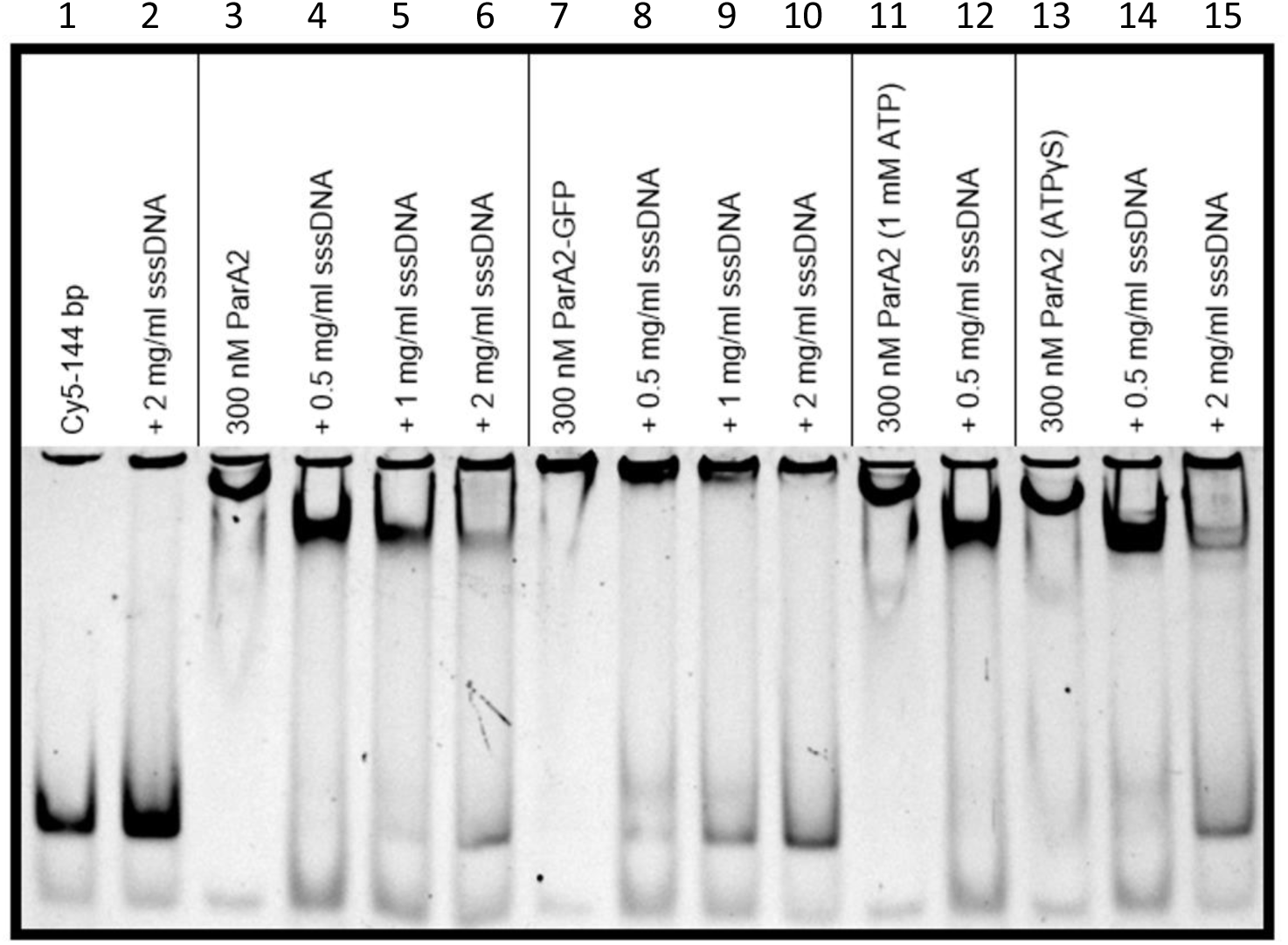
EMSA of ParA2 dissociation from DNA. ParA2-DNA complexes were preformed by incubation with 300 nM ParA2, 5 nM Cy5-labeled 144-bp DNA, 2 mM ATP (lanes 3, 7, 11, 13) unless stated otherwise. Controls of Cy3-labeled DNA and with sssDNA (lanes 1, 2) showed a dark band with fainter lower band at the bottom of gel, deemed as a nonspecific product from PCR. Increasing concentrations of sssDNA added (lanes 4-6) showed competition with bound DNA substrate (upper band) and increasing dissociation of complexes to free DNA (lower band). Complexes formed with ParA2-GFP-His (lanes 7-10) showed similar levels of dissociation as wt ParA2. Lower ATP concentrations of 1 mM (lanes 11, 12) showed similar dissociation as higher ATP concentrations (lanes 3, 4). Reactions incubated with 1 mM ATPγS (lanes 13-15) showed similar levels of DNA dissociation as for complexes with ATP.

**Figure S6.**
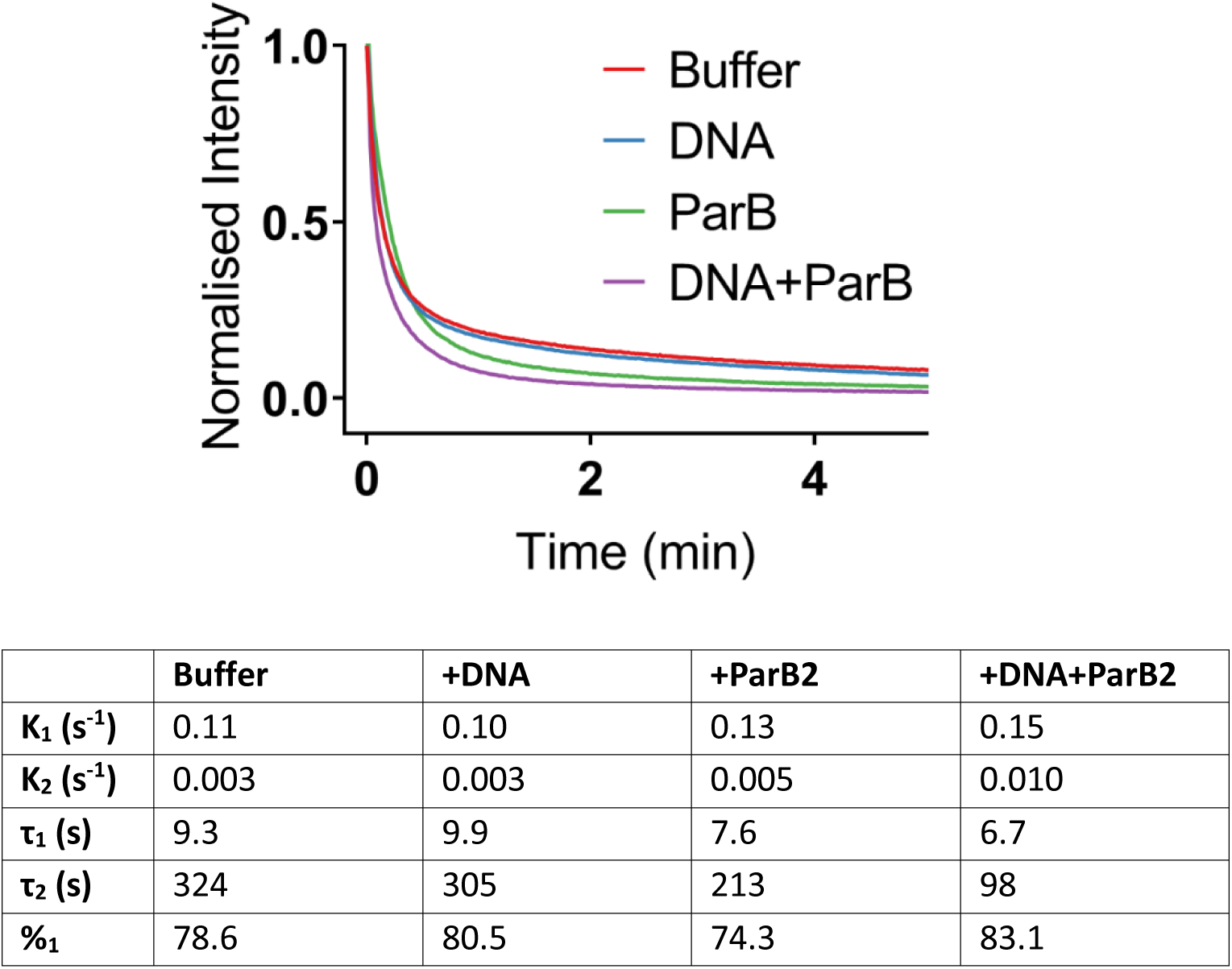
Dissociation of ParA2-GFP from DNA carpet with additional cofactors. Buffer containing 1 μM ParA2-GFP preincubated with ATP was infused into a DNA carpeted flowcell until steady state. Buffer flow was switched at t=0 to a wash buffer alone, or with 100 μg/ml DNA, 2 μM ParB2 or 100 μg/ml DNA and 2 μM ParB2. Fluorescence intensity was measured over time. Fluorescence intensities were subtracted for background and normalized to their prewash levels. The dissociation curves were fitted to a two-exponential decay model using GraphPad Prism 8.3 software. The presence of ParB2 increased the dissociation rate of ParA2-GFP from the DNA carpet. The addition of both ParB2 and DNA further increased the rate of ParA2-GFP dissociation.

**Figure S7.**
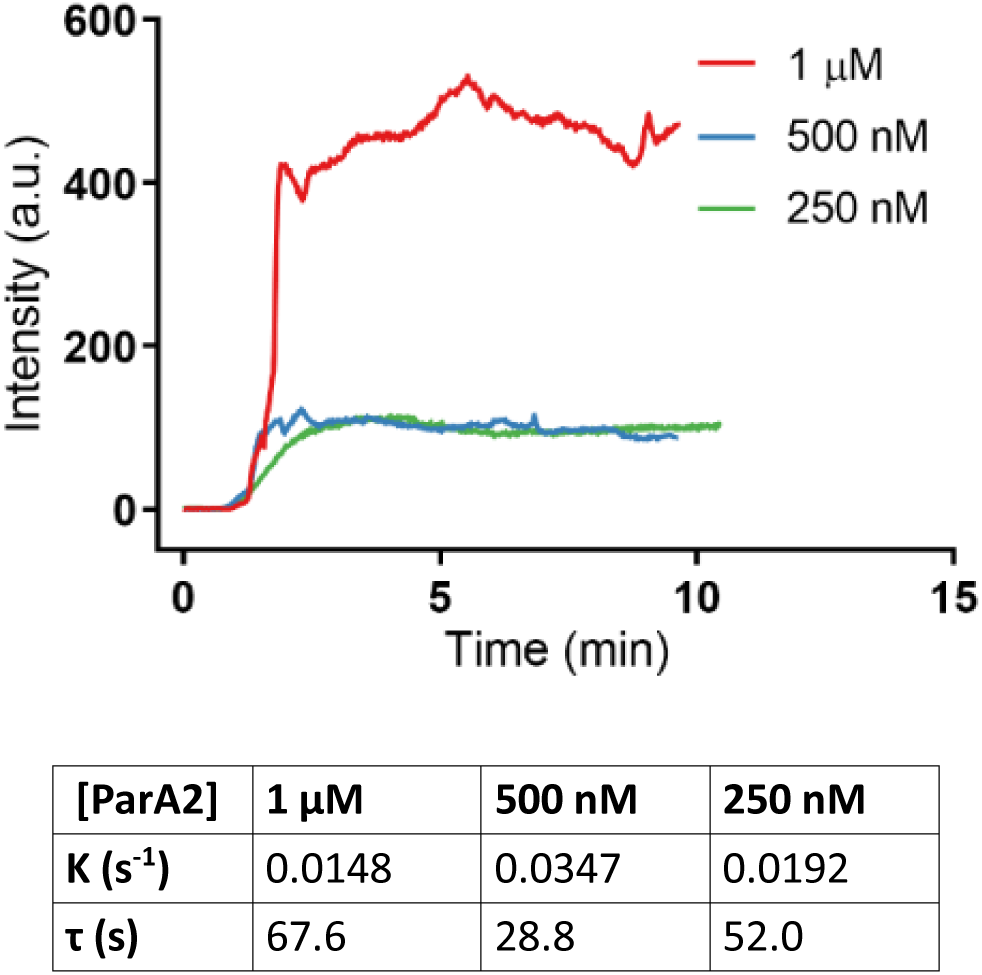
ATP start assay of ParA2-GFP. Separate solutions of 2, 1, 0.5 µM ParA2-GFP and 2 mM ATP were prepared in Par Buffer. The two solutions were loaded into 1 ml syringes and attached to a micro-static T-mixer (Upchurch). The T-mixer was attached to the single inlet port of a DNA-carpeted flow cell. Both samples were infused at 10 µl/min each with a combined flowrate of 20 µl/min entering the flowcell. Final concentrations of ParA2-GFP after mixing are as indicated. The flowcell was imaged with TIRFM close to the inlet port to minimize the time between the point of solution mixing and protein binding. Fluorescence intensity of ParA2-GFP plotted vs. time of sample infusion showed a lag time of ParA2-GFP binding the DNA carpet and lower intensities compared to ParA2-GFP preincubated with ATP. Binding curves were fitted to single exponential increase. Table shows fitted time constants and observed rates of DNA binding for each protein concentration.

**Table S1.** Kinetic rates of ParA2 interactions with adenine nucleotides.

| **ParA2** | **0.6125 μM** | **1.25 μM** | **2.5 μM** |
| --- | --- | --- | --- |
| **<u>ATP association</u>** |  |  |  |
| **K ($s^{-1}$)** | 0.132 ± 0.031 | 0.093 ± 0.013 | 0.093 ± 0.005 |
| **$\tau$ (s)** | 7.8 ± 1.8 | 11.7 ± 0.8 | 10.7 ± 0.6 |
| **<u>ATP dissociation</u>** |  |  |  |
| **K<sub>1</sub> ($s^{-1}$)** | 0.019 ± 0.001 | 0.019 ± 0.003 | 0.015 ± 0.002 |
| **$\tau_1$ (s)** | 51.0 ± 2.3 | 52.4 ± 0.9 | 60.6 ± 6.7 |
| **<u>ADP association</u>** |  |  |  |
| **K ($s^{-1}$)** | 0.162 ± 0.024 | 0.140 ± 0.008 | 0.121 ± 0.004 |
| **$\tau$ (s)** | 6.2 ± 0.9 | 7.2 ± 0.4 | 8.3 ± 0.3 |
| **<u>ADP dissociation</u>** |  |  |  |
| **K ($s^{-1}$)** | 0.072 ± 0.002 | 0.081 ± 0.004 | 0.081 ± 0.002 |
| **$\tau$ (s)** | 13.8 ± 0.4 | 12.4 ± 0.7 | 12.3 ± 0.3 |
| **<u>ADP → ATP</u>** |  |  |  |
| **K ($s^{-1}$)** | 0.023 ± 0.004 | 0.034 ± 0.001 | 0.054 ± 0.002 |
| **$\tau$ (s)** | 43.7 ± 7.8 | 28.2 ± 0.7 | 18.7 ± 0.8 |
| **<u>ADP → ADP</u>** |  |  |  |
| **K ($s^{-1}$)** | 0.068 ± 0.006 | 0.078 ± 0.005 | 0.097 ± 0.019 |
| **$\tau$ (s)** | 14.8 ± 1.3 | 12.9 ± 0.8 | 10.6 ± 2.3 |
| **<u>ATP → ATP</u>** |  |  |  |
| **K ($s^{-1}$)** | 0.026 ± 0.001 | 0.047 ± 0.008 | 0.092 ± 0.012 |
| **$\tau$ (s)** | 38.2 ± 0.8 | 21.5 ± 3.5 | 10.9 ± 1.4 |

Errors are ± SD

**Table S2.** ParA2 conformational change rates in the presence of ATP.

| **ParA2** | **0.6125 μM** | **1.25 μM** | **2.5 μM** |
| --- | --- | --- | --- |
| **<u>ParA2</u>** |  |  |  |
| **K ($s^{-1}$)** | 0.0177 ± 0.0016 | 0.0147 ± 0.0006 | 0.0156 ± 0.0015 |
| **$\tau$ (s)** | 56.76 ± 5.25 | 68.20 ± 2.75 | 64.22 ± 6.20 |
| **<u>ParA2 + DNA</u>** |  |  |  |
| **K ($s^{-1}$)** | 0.0328 ± 0.0015 | 0.0670 ± 0.0007 | 0.0684 ± 0.0006 |
| **$\tau$ (s)** | 30.53 ± 1.43 | 14.94 ± 0.16 | 14.63 ± 0.13 |
| **<u>ParA2 + ParB2</u>** |  |  |  |
| **K ($s^{-1}$)** | 0.0191 ± 0.0009 | 0.0149 ± 0.0001 | 0.0150 ± 0.0019 |
| **T (s)** | 52.43 ± 2.55 | 66.70 ± 0.23 | 67.17 ± 8.54 |
| **<u>ParA2 + ParB2 + DNA</u>** |  |  |  |
| **K ($s^{-1}$)** | 0.0361 ± 0.0015 | 0.0613 ± 0.0011 | 0.0702 ± 0.0007 |
| **$\tau$ (s)** | 27.69 ± 1.12 | 16.32 ± 0.30 | 14.31 ± 0.08 |

Errors are ± SD

**Table S3.** Binding and dissociation rates of ParA2-GFP on DNA carpet.

| ParA2-GFP | 1 μM (ATP) | 500 nM (ATP) | 250 nM (ATP) | 1 μM (ATPγS) |
| --- | --- | --- | --- | --- |
| **<u>Binding</u>** |  |  |  |  |
| **K (s<sup>-1</sup>)** | 0.053 ± 0.006 | 0.051 ± 0.009 | 0.049 ± 0.011 | 0.016 ± 0.004 |
| **$\tau$ (s)** | 18.8 ± 1.9 | 20.0 ± 3.4 | 21.1 ± 4.4 | 64.5 ± 14.6 |
| **<u>Dissociation</u>** |  |  |  |  |
| **K<sub>1</sub> (s<sup>-1</sup>)** | 0.11 ± 0.04 | 0.17 ± 0.08 | 0.19 ± 0.12 | 0.015 ± 0.002 |
| **$\tau_1$ (s)** | 10.0 ± 4.2 | 7.2 ± 4.5 | 7.0 ± 4.3 | 70.2 ± 9.7 |
| **K<sub>2</sub> (s<sup>-1</sup>)** | 0.0046 ± 0.0001 | 0.0048 ± 0.0001 | 0.0041 ± 0.0001 | 0.0016 ± 0.0002 |
| **$\tau_2$ (s)** | 218 ± 6 | 208 ± 6 | 242 ± 6 | 627 ± 6 |
| **%<sub>1</sub>** | 78.6 ± 4.1 | 79.7 ± 5.0 | 83.2 ± 1.3 | 48.7 ± 3.4 |

**K<sub>1</sub>:** Rate constant of fast decay species.
**K<sub>2</sub>:** Rate constant of slow decay species.
**$\tau_1$ :** Time constant of fast decay species.
**$\tau_2$ :** Time constant of slow decay species.
**%<sub>1</sub>:** Fraction of decay accountable to faster species.
Errors are ± SD

**Table S4.** Fitted FRAP recovery time constants and fractions of ParA2-GFP on DNA carpet.

| | Density (%) | $\tau_1$ (s) | $\tau_2$ (s) | $F_1$ | $F_2$ | $F_{\text{immobile}}$ |
| --- | --- | --- | --- | --- | --- | --- |
| ParA | 28 | $2.3 \pm 0.4$ | $121 \pm 32$ | 0.64 | 0.23 | 0.13 |
| | 100 | $8.5 \pm 1.8$ | $181 \pm 16$ | 0.44 | 0.32 | 0.24 |
| ParA:ParB (1:0.5) | 34 | $6.2 \pm 0.5$ | $241 \pm 28$ | 0.62 | 0.21 | 0.17 |
| ParA:ParB (1:1) | 45 | $6.6 \pm 1.5$ | $274 \pm 78$ | 0.47 | 0.18 | 0.38 |
| ParA:ParB (1:2) | 43 | $8.2 \pm 1.7$ | $308 \pm 81$ | 0.48 | 0.18 | 0.34 |

Means are an average of three FRAP measurements and errors are ± SD
**Density:** Percentage of protein coverage relative to intensity of 1 μM ParA2-GFP on DNA carpet at steady state.
$\tau_1$ : Photobleaching recovery time constant of fast species
$\tau_2$ : Photobleaching recovery time constant of slow species
$F_1$ : Fraction of fast species
$F_2$ : Fraction of slow species
$F_{\text{immobile}}$ : Fraction of immobile species that did not recover after post-bleach, $1-(F_1 + F_2)$
Errors are ± SD

## Materials and Methods

### ParA2 protein expression and purification

10 ml of LB medium with 50 µg/ml kanamycin (Kan) was inoculated with *E. coli* BL21(DE3) transformed with pSC01, and grown at 37°C overnight. Culture was added to 2×500 ml fresh LB/Kan to an OD_600_ 0.55. Cultures were cooled to 25°C and expression was induced with 1 mM isopropyl β-D-thiogalactoside (IPTG) for 2 h at 25°C. Cells were harvested by centrifugation for 20 min at 4,000 g and stored at −80°C. Cells obtained from a 1 L culture were defrosted and suspended in 10 ml per gram of pellet of 50 mM Tris-HCl pH 8.0, 0.1 M NaCl, along with ½ protease inhibitor tablet (Roche) and 1 mg/ml of lysozyme. Cells were placed on ice and disrupted by sonication using medium probe on a Soniprep 150 using 3 cycles of 20 s at 16-micron amplitude. Cell debris was removed by centrifugation at 72,000 g for 10 minutes at 24,500 rpm and the supernatant fraction was used for purification. ParA2 purification was performed on an FPLC AKTA system (GE Healthcare). The cell extract was applied on a 5 ml Heparin-HP cartridge (GE Healthcare) equilibrated in Tris buffer. Protein sample was eluted by a 50 ml gradient of 0−0.5 M NaCl in Tris buffer and 2.5 ml fractions were collected. The main peak containing protein was eluted at 0.25 M NaCl and 3-4 peak fractions were combined for further purification by anion exchange chromatography and gel filtration. The protein sample was diluted 2.5 fold with water to 0.1 M NaCl and applied on a 6 ml Resource Q column (GE Healthcare) equilibrated with Tris buffer. Elution was performed at 6 ml/min with 60 ml gradient of 0.1−0.7 M NaCl and 2.5 ml fractions collected. ParA2 was eluted at 0.35 M NaCl. Two peak fractions were combined and concentrated to 1 ml (Vivaspin 50,000 MWCO) and loaded on a 1.6×60 HiLoad Superdex 200 column equilibrated in 0.5 M NaCl 50mM Tris-HCl pH 8.0. Gel filtration was performed at 1.5 ml/min flow rate. Peak fractions were combined and concentrated. 2 mM DTT, 0.1 mM EDTA and 10 % glycerol were added, before storage at −80°C until further use. SDS-PAGE showed ParA2 yields with 98% purity. Protein sequence of ParA2 was analyzed with mass spectrometry.

### ParA2-GFP purification

10 ml of LB medium with 50 µg/ml ampicillin (Amp) was inoculated with *E. coli* BL21(DE3) transformed with pLCH02, and grown at 37°C overnight. Culture was added to 2 x 500 ml fresh LB/Amp to an OD_600_ 0.5. Cultures were cooled to 16°C and expression was induced with 1 mM isopropyl β-D-thiogalactoside (IPTG) overnight at 16°C. Cells were harvested by centrifugation for 20 min at 4,000x*g* and stored at −80°C. Cells obtained from a 1 L culture were defrosted and suspended in 10 ml per gram of pellet of 50 mM Tris-HCl pH 8.0, 0.1 M NaCl, along with ½ a protease inhibitor tablet (Roche) and 1 mg/ml of lysozyme. Cells were placed on ice and disrupted by sonication on a Soniprep 150 machine. Three cycles of 20 s treatment at 16-micron amplitude were applied with cooling between the treatments. Cell debris was removed by centrifugation at 72,000x*g* for 10 min at 24,500 rpm. The supernatant fraction (cell free extract, CFE) was separated and used for purification. CFE was applied on a 5 ml His-Trap HP column (GE Healthcare) equilibrated in 50 mM Tris-HCl pH 8.0, 0.1 M NaCl, with an AKTA purifier system with a flow rate of 5ml/min. Bound protein was eluted by a 50 ml gradient of imidazole from 0 to 0.5 M in 50 mM Tris-HCl pH 8.0, 0.1 M NaCl. Peak fractions were combined, and the volume of the protein sample was reduced to <2 ml. Sample was applied to 1.6 x 60 ml HiLoad Superdex 200 column equilibrated in buffer A. Gel filtration was performed at 1.5 ml/min flow rate. 2 ml fractions were collected after void volume. Peak fractions were combined and concentrated. 0.3 mg of TEV protease was added per 1 mg ParA2-GFP-His in 50 mM Tris pH 8.0, 150 mM NaCl and left overnight at 16°C. Sample was suspended in 25 mM Tris pH 8.0, 500 mM NaCl, 20 mM imidazole, 10% glycerol and 2 mM BME and loaded onto a 5 ml HisTrap column, eluting over a 12 CV gradient. The flow through was collected and reloaded onto the column to run once more. Protein was eluted with 25 mM Tris pH 8.0, 500 mM NaCl, 1 M imidazole, 10% glycerol and 2 mM BME. Peak fractions were collected, concentrated to <2 ml and buffer exchanged into 50 mM Tris pH 8.0, 500 mM NaCl, 10% glycerol, 2 mM DTT, and 0.1 mM EDTA using a VivaSpin column. The sample was then loaded onto a Superdex 200 16/600 and eluted over a 1.2 CV isocratic gradient. Peak fractions were pooled, concentrated, and stored at −80°C.

### ParB2 purification

5 ml of LB medium with 100 µg/ml ampicilin (Amp) was inoculated with *E. coli* BL21(DE3) transformed with pLCH04 and grown as in section 2.2.4. Culture was added to 500 ml fresh LB/Amp to an OD600 0.55. Cultures were cooled to 25 °C and expression was induced with 1 mM IPTG for 4 h at 25 °C. Cells were harvested as described for ParA2, in section 2.2.4. CFE was applied on a 5 ml His-Trap HP column (GE Healthcare) equilibrated in 50 mM Tris-HCl pH 8.0, 0.1 M NaCl at flow rate 5ml/min. Bound protein was eluted by a 50 ml gradient of imidazole from 0 to 0.35 M in 50 mM Tris-HCl pH 8.0, 0.1 M NaCl. Peak fractions were combined, and the volume of the protein sample was reduced to <2 ml (Vivaspin 50,000 MWCO). Sample was applied to 1.6 x 60 ml HiLoad Superdex 200 column equilibrated in 0.5 M NaCl 50mM Tris-HCl pH 8.0. Gel filtration was performed at 1.5 ml/min flow rate and 2 ml fractions were collected after void volume. Peak fractions were combined and concentrated (Vivaspin 50,000 MWCO). The TEV-cleavage protocol for ParB2-His was performed as for ParA2-GFP-His above. Protein was concentrated, and stored at −80 °C

### TIRF microscopy

A home-built prism-based TIRFM was set up using a Ti-Eclipse (Nikon) microscope with a PlanApo 100x NA 1.45 oil-immersion objectives. Laser excitation light of 488 nm (Cobolt 06-MLD) at laser power 100 μW was focused and aligned onto a prism placed on a quartz slide for TIRF illumination at the center of the objectives. The fluorescence emission light was filtered by a notch filter (Thorlabs NF488-15) to remove the excitation laser line and an emission bandpass filter (Chroma ET535/70m). Fluorescence images were captured using a sCMOS camera (Prime 95B, Photometrics) with exposure time 100 ms, frame rate 1 s at 16-bit depth. The camera bias of 100 arbitrary units was subtracted from measured intensity. Micro-manager software was used for camera control and image acquisition. ImageJ (NIH) was used for image analysis. Syringe pumps (WPI) were controlled using RealTerm open software.

### FRAP experiments

A free-space 488 nm laser (Coherent) was aligned to the TIRF microscope via the backport for photobleaching. The laser beam was focused by a meniscus lens (f=-500 mm, Thorlabs) and reflected by a ZT488/640rpc dichroic mirror (Chroma) into the back of the objectives. ParA2-GFP was preincubated at 10 μM in the presence of 2 mM ATP for 15 minutes at 25°C and diluted to 1 μM ParA2-GFP for infusion to various densities in the flowcell. Camera acquisition was at 1 frame/s and exposure time 100 ms. Acquisition was stopped for ∼1 s as the DNA carpet was photobleached to around 50% of initial intensity, before continuing acquisition. FRAP curves were normalized and corrected for background intensity and overall background bleaching due to FRAP. Each curve was fitted to a double exponential equation in Origin Pro software.

### Construction of strains for microscopy

A plasmid expressing ParA2-fused to GFP was electroporated into CVC209 (see Strain list) and used for microscopy as described below. First, the *parA2:gfp* DNA fragment was amplified from pLCH02 template DNA using primers LCH11-A2gfp-fwd and LCH12-A2gfp-rev. The DNA fragment was then inserted into pBAD-HisB to construct plasmid pLCH08. The ampicillin resistance gene cassette was interrupted by inserting a *kanr*2 DNA fragment (amplified from pACYC177) thus conferring kanamycin resistance and resulting in plasmid pLCH10.

### Strain List

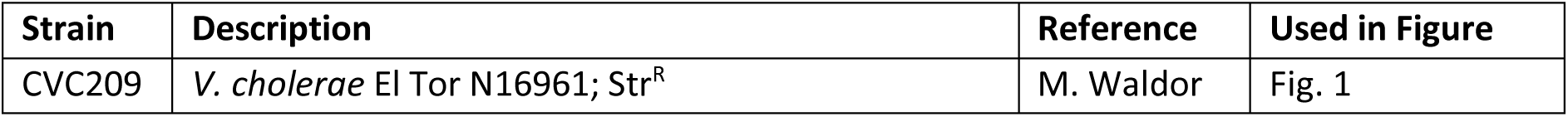

### Plasmid List

Plasmid List:
| Plasmid | Description | Reference | Used in Figure |
| --- | --- | --- | --- |
| pLCH02 | pET15b-*parA2*-GFP-*his*; Ap<sup>R</sup> | This Study | Figures 1, 6 |
| pLCH08 | P<sub>BAD</sub>-*parA2:GFP* in pBAD-HisB; pBR ori; Ap<sup>R</sup> | This Study | Figure 1 |
| pLCH10 | P<sub>BAD</sub>-*parA2:GFP* in pBAD-HisB; pBR ori; Kn<sup>R</sup> | This Study | Figure 1 |
| pSC01 | *parA2* in pET28 b (+); Kn<sup>R</sup> | This Study | Figures 2-5 |
| pLCH04 | *parB2-his* in pET15 b (+); Ap<sup>R</sup> | This Study | Figures 2, 4, 6 |
| pBKSII | pBluescriptII KS (+) | Stratagene | Figure 5 |

### Primer List

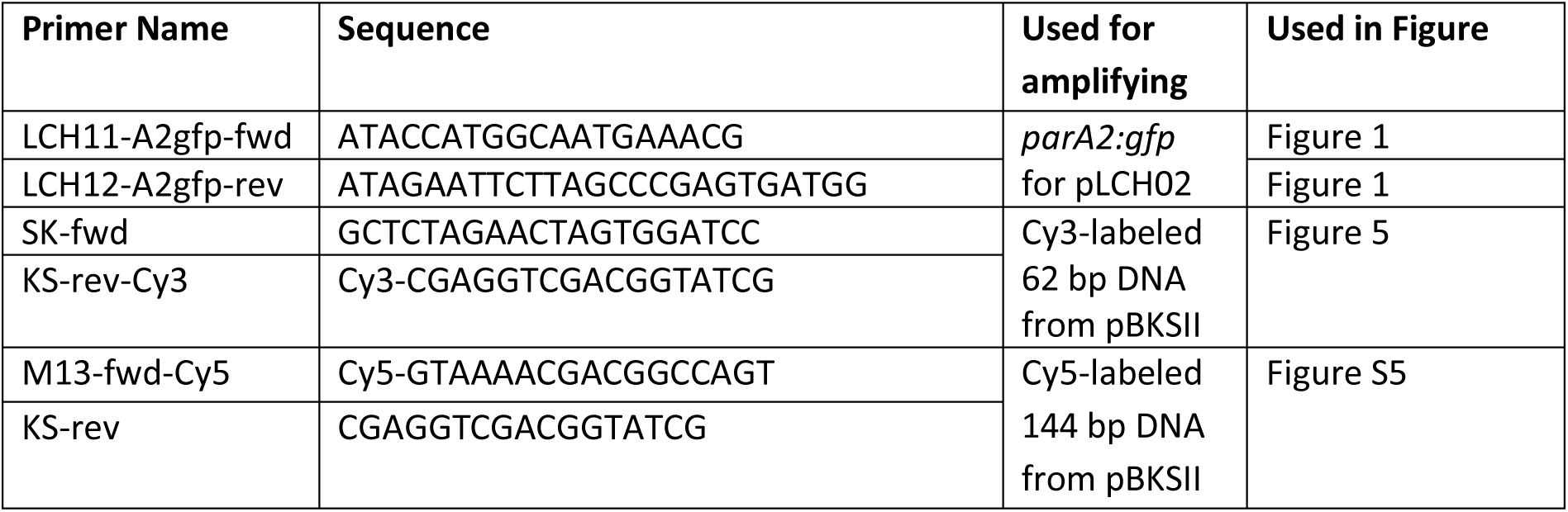

